# Dietary and serum tyrosine, white matter microstructure and inter-individual variability in executive functions in overweight adults: relation to sex/gender and age

**DOI:** 10.1101/2021.12.01.470815

**Authors:** Brecht A-K, E Medawar, R Thieleking, J Sacher, F Beyer, A Villringer, AV Witte

## Abstract

Tyrosine (tyr), the precursor of the neurotransmitter dopamine, is known to modulate cognitive functions including executive attention. Tyr supplementation is suggested to influence dopamine-modulated cognitive performance. However, results are inconclusive, regarding the presence or strength and also the direction of the association between tyr and cognitive function. This pre-registered cross-sectional analysis investigates whether diet-associated serum tyr relates to executive attention performance, and whether this relationship is moderated by differences in white matter microstructure. 59 healthy, overweight, young to middle-aged adults (20F, 28.3 ± 6.6 years, BMI: 27.3 ± 1.5 kg/m^2^) drawn from a longitudinal study reported dietary habits, donated blood and completed diffusion-weighted brain magnetic resonance imaging and the attention network test. Main analyses were performed using linear regressions and non-parametric voxel-wise inference testing.

Confirmatory analyses did neither support an association between dietary and serum tyr nor a relationship between relative serum tyr/large neutral amino acids (LNAA) levels or white matter microstructure and executive attention performance. However, exploratory analyses revealed higher tyr intake, higher serum tyr and better executive attention performance in the male sex/gender group. In addition, older age was associated with higher dietary tyr intake and lower fractional anisotropy in a widespread cluster across the brain. Finally, a positive association between relative serum tyr/LNAA and executive attention performance was found in the male sex/gender group when accounting for age effects.

Our analysis advances the field of dopamine-modulated cognitive functions by revealing sex/gender and age differences which might be diet-related. Longitudinal or intervention studies and larger sample sizes are needed to provide more reliable evidence for links between tyr and executive attention.

## Introduction

Dietary intake of the amino acid tyrosine (tyr), a precursor of the neurotransmitter dopamine, has been claimed to modulate brain functions and cognitive performance, such as executive functions (Aquili, 2020). In rodents, protein-rich diets (single meals or habitual intake) or direct tyr injection increase serum and brain tyr concentrations and dopamine (DA) synthesis in the brain (reviewed in (John D. Fernstrom & Fernstrom, 2007)). Human studies also showed that dietary tyr/protein intake changes serum levels of tyr (J D Fernstrom et al., 1979; Strang et al., 2017; van de Rest et al., 2017; Wurtman et al., 2003), and acute depletion of tyr decreased brain DA measured indirectly using positron emission tomography (PET) (Leyton et al., 2004). Considering cognitive effects, some studies showed that tyr intake correlated with differences in inhibitory control (Colzato et al., 2014), task switching (Steenbergen et al., 2015) as well as working memory performance (Colzato et al., 2013; Hensel et al., 2019; Kühn et al., 2019; Thomas et al., 1999; van de Rest et al., 2017) and reward processing (Aquili, 2020), resulting in either improved (habitual dietary tyr or protein shakes) or both improved and weakened (high-dose oral supplementary tyr) test performance. In parallel, other studies could not demonstrate significant effects of tyr intake on executive function such as conflict monitoring and resolution measured with the attention network task (ANT) (Frings et al., 2020) or with a related task combining subliminal priming and flanker interference (Stock et al., 2018).

These at first glance contradictive findings might be insightful considering several important aspects of the tyr-DA and DA-cognition relationship: Firstly, the direction and strength of DA-enhancing drug effects on task performance has been found to depend on differences in baseline DA levels. This baseline dependence may be modelled by an inverted U-shaped dose-response curve, leading to either improvement, maintenance, or impairment of task performance by increasing DA availability (Cools & D’Esposito, 2011). The inverted U-shaped curve presumably relates to complex autoregulatory mechanisms of the dopaminergic system which controls DA synthesis and release (Cools, 2019), including tyr concentration-dependent activity differences of brain tyr hydroxylase (Reed et al., 2010). Secondly, the ratio of how much serum tyr reaches the brain depends on blood concentrations of other large neutral amino acids (LNAA; i.e. tryptophan, phenylalanine, leucine, isoleucine, valine, methionine) (Wurtman et al., 2003) due to a competitive transport system at the carrier site (J. D. Fernstrom, 1983). Therefore, it is crucial (1) to consider baseline DA levels and (2) to measure tyr/LNAAs ratios in blood to estimate tyr levels in the brain (John D. Fernstrom & Fernstrom, 2007). Notably, blood LNAAs also change in response to insulin secretion and related uptake of amino acids into peripheral tissues (John D. Fernstrom, 2013), therefore overall macronutrient composition, intake and diurnal changes in the metabolism need to be considered when evaluating how dietary tyr affects DA-related cognitive function (J D Fernstrom et al., 1979; Strang et al., 2017; Wurtman et al., 2003).

In addition, dopaminergic drug effects have been suggested to also depend on inter-individual differences in microstructural properties of white matter tracts that are associated with targeted cognitive function (van der Schaaf et al., 2013; Martine R. Van Schouwenburg et al., 2013). The relevance of white matter connectivity is also supported by numerous studies indicating structure-function associations (Chaddock-Heyman et al., 2013; Cremers et al., 2016; Niogi et al., 2010; M. R. Van Schouwenburg et al., 2014; Vinçon-Leite et al., 2020; Yin et al., 2013). Therefore, differences in regional white matter microstructure should be considered when examining the effects of tyr on cognitive performance.

Taken together, the relationship between dietary tyr intake and cognitive performance is suggested rather complex and requires a careful consideration of numerous interacting mechanisms and conditions which have not been fully addressed in studies on executive function. We therefore aimed (1) to examine how inter-individual differences in self-reported dietary tyr intake relate to individual differences in serum tyr levels and (2) to determine whether differences in serum tyr (and tyr/LNAA ratio) could explain variation in executive attention performance, measured with the ANT in a sample of young to middle-aged adults with omnivorous, naive eating habits resembling a typical Western European diet. Moreover, we examined possible structure-function relationships to address whether the strength and direction of potential associations between serum tyr and executive attention performance might depend on differences in white matter microstructure of the executive attention network. To investigate possible structure-function relationships, we calculated tract-based fractional anisotropy (FA) based on diffusion weighted magnetic resonance imaging (dwMRI). We pre-registered all hypotheses and analyses at https://osf.io/hbjyr to increase transparency and reproducibility. A visualisation of our hypotheses is presented in Figure 1.

**Figure 1:**
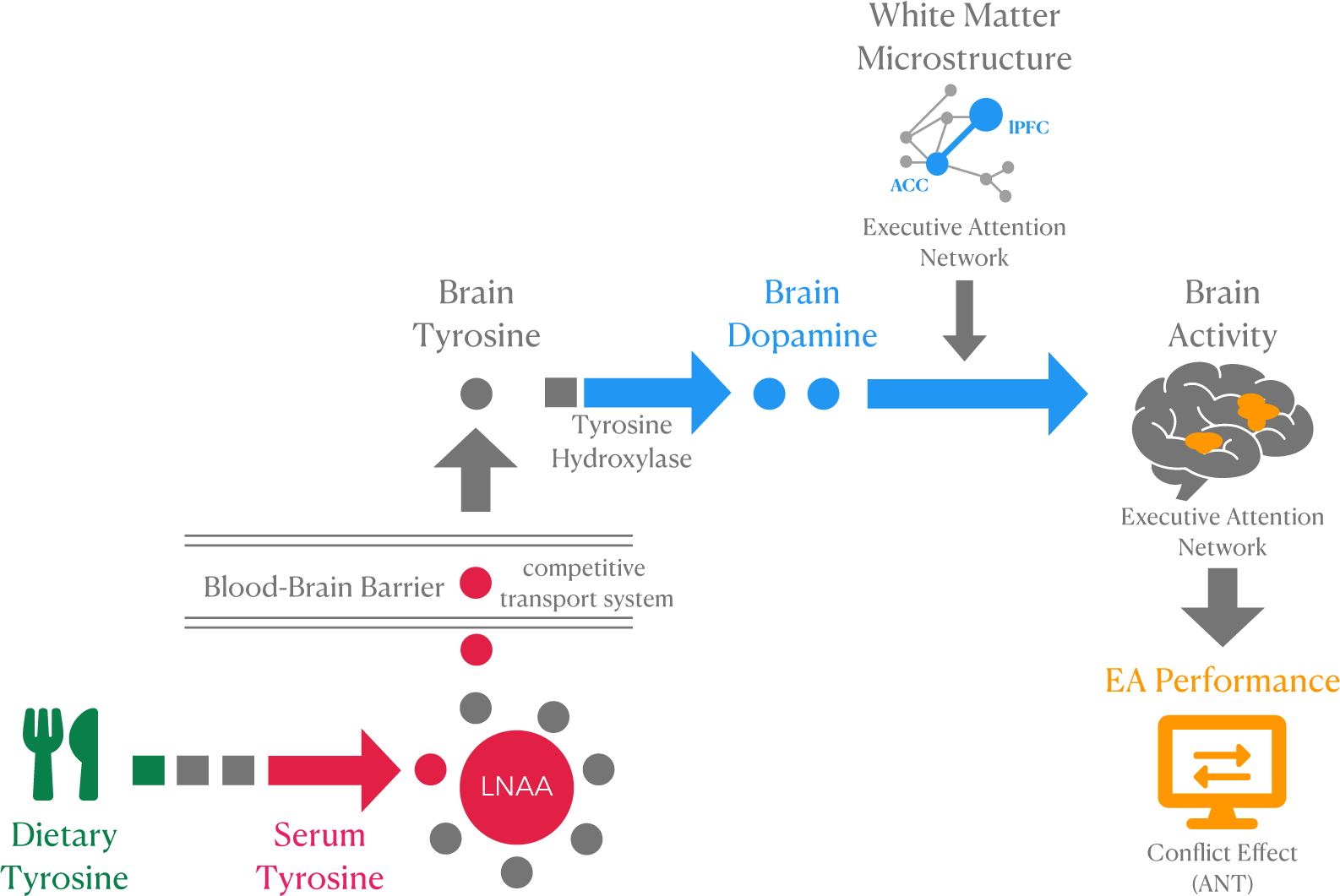
Hypothesized mechanism underlying a link between tyrosine intake and executive attention performance. Abbreviations: LNAA, large neutral amino acids; ACC, anterior cingulate cortex; lPFC, lateral prefrontal cortex; EA, executive attention; ANT, attention network test.

## Methods

### Study design and participants

Data was drawn from a two-arm cross-over within-subject randomized controlled study with four time points (https://clinicaltrials.gov/ct2/show/NCT03829189). In total, 60 young to middle-aged adults were recruited from the participant database of the Max Planck Institute for Human Cognitive and Brain Sciences (Leipzig, Germany) or via advertisements. The institutional ethics board of the Medical Faculty of the University of Leipzig, Germany, raised no concerns regarding the study protocol (228/18-ek) and all participants provided written informed consent. They received reimbursement for participation. For the current analysis, baseline assessments are drawn from the longitudinal data set.

Participants were included based on body mass index (BMI; range 25-30kg/m²) and eating behaviour, i.e. being on an omnivorous diet without food allergies or restrictions. All included subjects showed no unconventional eating habits or daily consumption of > 50g alcohol, >10 cigarettes, or > 6 cups of coffee. Exclusion criteria were neurological or psychiatric disorders, severe metabolic or internal disease or any medications acting on the central nervous system. Participants were excluded if they were pregnant or breastfeeding or females who were not using any hormonal contraceptive (pill, IUD or vaginal ring). Due to high depressive symptoms at testing day, one subject was excluded from data analysis. The final sample size comprises n=59 (female sex/gender= 20, male sex/gender = 39) healthy, overweight (BMI: 27.3 ± 1.5 kg/m^2^), young to middle-aged adults (28.3 ± 6.6 years, 19-45 years). Testing sessions were scheduled at either 07.15, 08.00, 09.15, 10.30 or 11.15 a.m. and began with the filling in of questionnaires followed by blood drawing, physiological measurements and MRI scanning. The cognitive assessment was conducted post-MRI 3 to 3.5 hours after the blood drawing.

### Attention Network Test

To measure individual differences in the efficiency of the executive attention network, we used a computerized version of the *Attention Network Test* (ANT) developed by Fan, Posner and colleagues (Fan et al., 2002). ANT data of n=58 was available; one data set was faulty due to technical problems. The participants’ task is to identify the direction of a centrally presented arrow flanked by neutral (lines), congruent (arrows in the same direction), or incongruent (arrows of opposite direction) stimuli (see Figure). In addition, the task includes four cue conditions (no cue, center cue, double cue, spatial cue) that precede the presentation of the stimuli and could indicate the location of the upcoming target stimuli (only spatial cue). A 30-minute test session consists of a practice block of 24 trials in which subjects receive feedback, and 3 experimental blocks consisting of 96 randomly ordered trials without feedback. The efficiency of each network is quantified by error rates (ER) and reaction times (RT). ER represent an accuracy score. RT of only correct trials are used to compute so-called alerting, orienting and conflict effects that quantify respective performance in one of the three attentional components. Conflict effects measure the processing of conflicting visual information (i.e., the influence of incongruent flankers on the processing of a target stimulus). The smaller the difference between the RT in processing incongruent and congruent visual information, the better the EA performance.

*Conflict effects* – as a measure for the efficiency of the EA network – were calculated by subtracting mean RT of all congruent from mean RT of all incongruent trials. In addition, *alerting effects* were calculated by subtracting the mean RT of all double cue trails from the mean RT of all no cue trails and *orienting effects* were calculated by subtracting the mean RT of all spatial cue trails from the mean RT of all center cue trails.

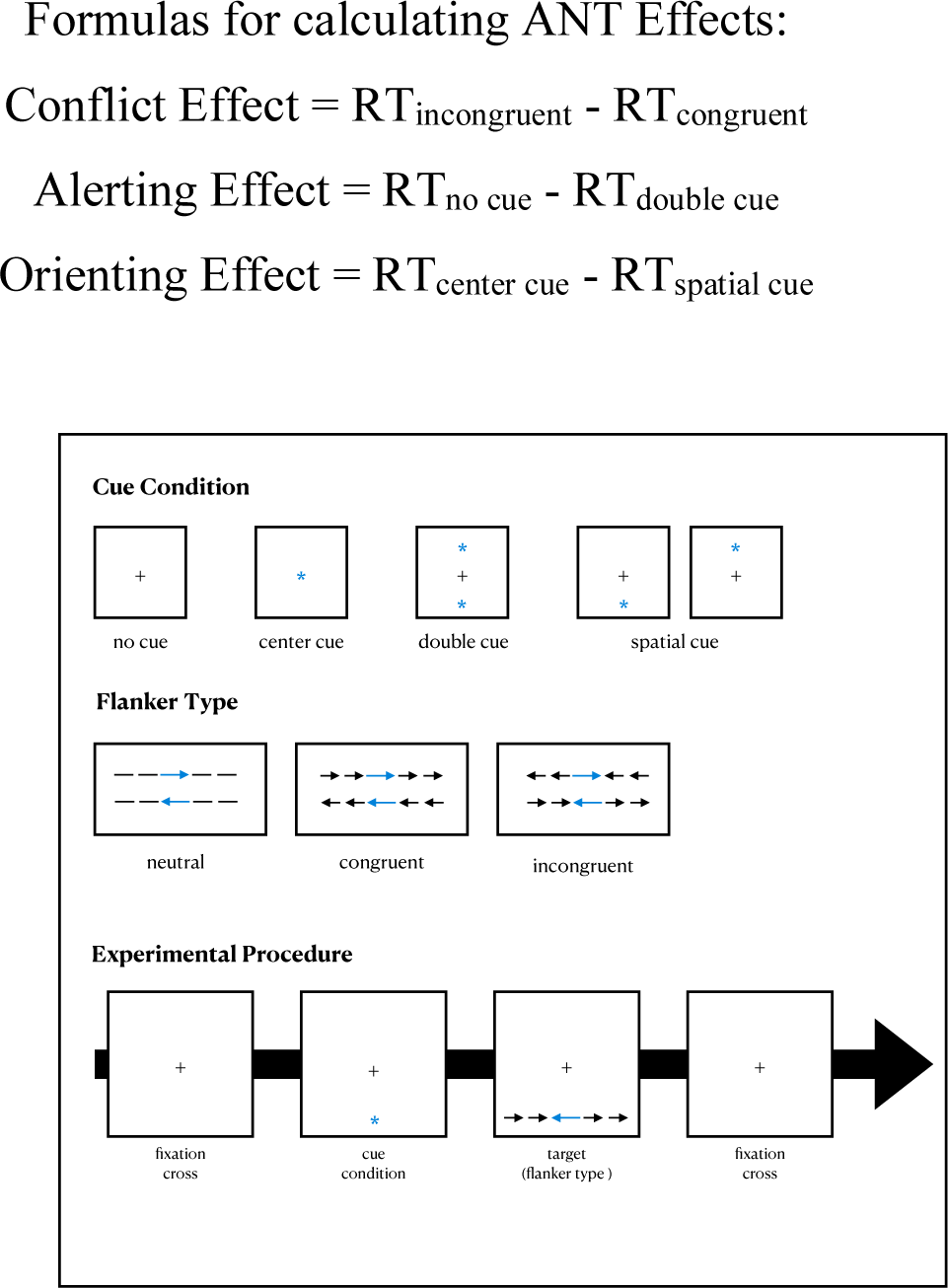

Experimental procedure of the Attention Network Test reproduced according to *Fan et al. (2002)*.

### Self-reported dietary tyrosine intake

Habitual dietary tyr intake over the last 7 days was assessed with the German Food Frequency Questionnaire (FFQ) DEGS1 by the Robert Koch Institute (Berlin, Germany); a tool to measure the intake of 53 single food items consumed based on self-reported frequency and quantity (Haftenberger et al., 2010). To get a measure of dietary tyr intake, we calculated the mean tyr intake per day using reference food items from the German Nutrient Database (Bundeslebensmittelschlüssel (BLS), Version 3.02) or - in rare cases - individual sources directly from food suppliers (e.g., for plant-based milk). Further details on nutrient scoring: https://osf.io/h73wj/. First, we controlled tyr intake for total kcal to put very high or low tyr intake into perspective. Second, we accounted for the intake of other dietary LNAAs to provide a more meaningful measure of dietary tyr intake and its potential effect in the brain (cf. competing transport system at the blood-brain barrier).

### Blood markers

Blood sampling was performed after overnight fasting (12.5 ± 2.2 h fasted) and about 2.5 hours before the ANT. The following amino acids were of interest: tyr, tryptophan, phenylalanine, leucine/isoleucine, valine, methionine. Blood samples were centrifuged and stored at -80 °C. Analyses were conducted at the Institute for Laboratory Medicine, Clinical Chemistry and Molecular Diagnostics (ILM) Leipzig University, Leipzig, Germany. Missing values due to limit of detection (LLOD) or limit of quantification (LLOQ) were substituted with values just below the limits (LLOD - 0.1 and LLOQ - 0.1). To relate dietary intake of tyr with serum tyr levels, absolute tyr levels were used. To get a proxy for acute brain DA levels, relative serum tyr/LNAAs ratios were calculated. Serum tyr/LNAAs ratios were calculated by dividing the tyr concentration by the summed concentration of all remaining LNAAs, i.e., tryptophan, phenylalanine, leucine, isoleucine, valine, methionine.

### Sex/Gender

Sex/gender variable was collected by asking participants in German about their “Geschlecht” with three response options: “*weiblich*”, “*männlich*” and “*keine Antwort”*. The third option was not chosen by anyone. The German word “Geschlecht” was used ambiguously, since the German term carries both the meaning of biological sex and social gender without further specification. This conceptual indeterminacy affects the two response options “weiblich” (female/feminine) and “männlich” (male/masculine).

Unfortunately, no other option such as “diverse” or “third gender” was provided. Since we cannot be certain which concept (i.e., biological sex or social gender) has been captured, we use the composite term *sex/gender* throughout the paper to highlight the underdetermination. Since a dichotomous question was asked, we will maintain this distinction between two groups by using the terms “*male sex/gender group”* and “*female sex/gender group”*. The reluctance to clearly assign the measured variable to biological sex or social gender will be particularly important when interpreting the results.

### Socio-economic status

Socio-economic status (SES) was included as a control variable because differences in both cognitive performance and tyr intake may be due to differences in SES (Cinelli et al., 2020; Kühn et al., 2019). SES was assessed using a questionnaire on SES and subjective social status (Lampert et al., 2013). The SES index was calculated as a sum value based on three sub-dimensions: education, occupation and income (theoretical range: 3 to 21 points). The SES index was considered as a continuous variable. The SES index was available for n=51 participants, eight participants did not provide information.

### Image acquisition and processing

*Anatomical MRI*. Anatomical MRI was acquired with a T1-weighted Magnetization Prepared - RApid Gradient Echo (MP-RAGE) sequence using the Alzheimer’s Disease Neuroimaging Initiative (ADNI) protocol http://adni.loni.usc.edu/methods/documents/mri-protocols/ with the following parameters: repetition time (TR) = 2300 ms; echo time (TE) = 2.98 ms; flip angle = 9°; Field-of-view (FOV): (256 mm)²; voxel size: (1.0 mm)³; 176 slices. Preprocessing included skull stripping and realignment to anterior and posterior commissure and tissue segmentation with the default settings using SPM12.

Diffusion-weighted MRI (dwMRI). dwMRI was acquired using the following parameters: TR = 5200 ms; TE = 75 ms; flip angle = 90°; FOV: (220 mm)²; voxel size: (1.7mm)³; 88 slices; max. b = 1000 s/mm² in 60 diffusion directions; partial Fourier = 7/8; GeneRalized Autocalibrating Partially Parallel Acquisitions (GRAPPA) factor = 2 (Griswold et al., 2002); interpolation = OFF. Ap/pa-encoded b0-images were acquired for distortion correction. We acquired n=58 dwMRI data sets; one participant aborted the MRI session due to nausea before dwMRI acquisition.

Preprocessing was performed with standard pipelines, including denoising (MRtrix v3.0) of the raw data, removal of gibbs!ringing artifact from all b0 images using the local subvoxel-shift method (Kellner et al., 2016) and outlier replacement using the eddy tool in FMRIB Software Library (FSL) (Andersson & Sotiropoulos, 2016). Subsequently, data was corrected for head motion and linearly co-registered to the T1 image with *Lipsia tools* (Lohmann et al., 2001). Finally, we applied tensor model fitting and generated fractional anisotropy (FA) images as an index of white matter coherence.

Through tract-based spatial statistics (TBSS) (Smith et al., 2006), we obtained FA maps of the individual white matter skeletons and extracted the mean FA value for each participant. All FA maps were co-registered using affine and non-linear transformations to FMRIB58_FA standard space and the individual local maximal FA values were projected onto the standard FA skeleton to match individual’s anatomy. The threshold for these standardized white matter fiber tract maps was set at 0.2. The FA skeleton maps were fed into voxel-wise analysis of FA for statistical comparison using the randomise tool by FSL version 5.0.11. We used 10,000 permutations (Winkler et al., 2014) in the confirmatory analyses and 5,000 permutations in the exploratory analyses concerning age correlations. Threshold-free cluster enhancement (TFCE) was applied as test statistic (Smith & Nichols, 2009) and significance level was set at α(FWE)=0.05. Voxel-wise analysis was conducted on a whole-brain level and in the exploratory age correlation analysis we controlled for frame-wise displacement as a measure of head motion (Beyer et al., 2017).

### Statistical analyses

Based on the OSF-preregistration, we distinguished between *pre-registered confirmator*y (systematically testing four hypotheses) and pre-registered *sensitivity* or *exploratory* analyses (including additional control variables). In addition, we performed additional *non-pre-registered exploratory* analyses to further elucidate possible relationships or underlying mechanisms. The selection and definition of the non-registered analyses builds on the results of the pre-registered analyses. All statistical analyses (except for brain data) were performed in *R* (version 4.0.3). Regression models and prerequisite testing were computed with *R base* functions, plots were created using *R base* functions or the R package *ggplot2*. The level of significance was set at α = 0.05.

#### (1) Dietary - Serum Tyrosine Relationship (pre-registered)

To investigate the relationship of dietary tyr intake and absolute serum tyr levels, we performed a multiple linear regression analysis controlling for sex/gender.

**Regression model:**

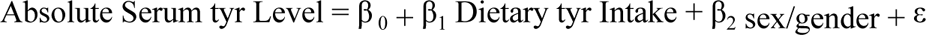

#### (2) Serum Tyrosine and Executive Attention (pre-registered)

To examine the relationship between serum tyr/LNAAs ratio and executive attention performance (i.e., conflict effects), we performed multiple linear regression analyses controlling for sex/gender. Executive attention performance was quantified by calculating conflict effects based on reaction times (RT). The serum tyr/LNAAs ratio was calculated by dividing the serum tyr level by the sum of the remaining LNAA.

**Regression model:**

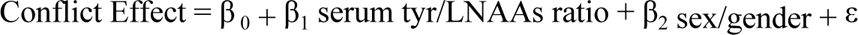

#### (3) Executive Attention and White Matter Microstructure (pre-registered)

To examine the neural basis of inter-individual variability in executive attention performance, we performed voxel-wise cross-subject whole-brain analyses on the FA skeletons to correlate *fractional anisotropy* (FA) values with conflict effects.

**General Linear Model:**

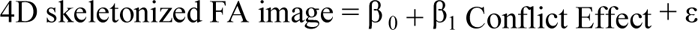

#### (4) Moderation Analysis (pre-registered)

Because of the lack of significant correlations between executive attention performance and FA values during the aforementioned voxel-wise whole-brain correlation analysis, this moderation analysis was not conducted.

### Sensitivity and Exploratory Analyses

In subsequent sensitivity analyses, additional control variables (BMI, SES, age) were added (Table 1 & **Fehler! Verweisquelle konnte nicht gefunden werden.**). We applied a *hierarchical regression analysis approach* (Lewis, 2007), adding each variable one after the other to examine the contribution of each predictor to the model. The level of significance was set at α = 0.05 for all sensitivity analyses.

**Table 1:** Sensitivity Analyses Models: Executive Attention and Serum tyr

| **Models: Executive Attention and Serum tyr** |
| --- |
| Conflict Effect = $\beta_0 + \beta_1$ serum tyr/LNAAs ratio + $\beta_2$ sex/gender + $\beta_3$ age + $\varepsilon$ |
| Conflict Effect = $\beta_0 + \beta_1$ serum tyr/LNAAs ratio + $\beta_2$ sex/gender + $\beta_3$ age + $\beta_4$ BMI + $\varepsilon$ |
| Conflict Effect = $\beta_0 + \beta_1$ serum tyr/LNAAs ratio + $\beta_2$ sex/gender + $\beta_3$ age + $\beta_4$ BMI + $\beta_5$ SES + $\varepsilon$ |

In addition, the multiple regression analysis was repeated for absolute dietary tyr intake as predictor variables to examine which tyr measures (self-reported dietary tyr intake vs. tyr serum level) better predicted differences in executive attention performance. Following Hensel et al. and Kühn et al., we adjusted dietary tyr intake to body weight. In addition, total energy intake was included as control variable.

**Regression Model:**

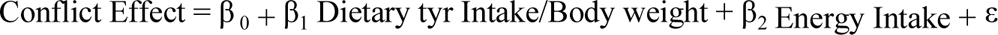

In addition, for the ANT, a two-way Analysis of Variance (ANOVA) was calculated to examine the influence of the two factors (1) flanker type (3 factor levels: neutral, congruent, incongruent) and (2) cue condition (4 factor levels: no cue, central cue, double cue, spatial cue) on the measured continuous dependent variable reaction times (RT) and error rates (ER), respectively. ANOVAs, post-hoc analyses for multiple pairwise comparisons (i.e., Tukey’s test), and Shapiro-Wilk normality test were calculated using R base functions. Homoscedasticity was tested with Levene’s test using the R package car.

### Non-Pre-registered Analyses

First, we explored associations between dietary tyr and serum tyr markers using Pearson’s correlation coefficients for the following relationships:

- Dietary tyr intake – serum tyr levels
- Dietary tyr intake – serum tyr/LNAAs ratios
- Dietary tyr intake (adjusted for energy intake) – serum tyr levels
- Dietary tyr intake (adjusted for body weight) – serum tyr levels
- Absolute serum tyr levels - relative serum tyr/LNAAs ratios

Correlational models (*Pearson’s product-moment correlation*) and prerequisite testing were computed with R base functions. Next, we examined sex/gender differences in variables of interest using non-parametric Mann-Whitney U test (i.e., Wilcoxon rank sum test), i.e.: dietary tyr intake, dietary tyr intake adjusted for energy intake, absolute serum tyr levels, relative tyr tyr ratios, ANT conflict effects RT, age, BMI, SES. Analyses were performed using the R package stats. The level of significance was set at α = 0.05. In addition, we performed the regression analyses outlined above with serum tyr/LNAAs ratio as predictor variable and executive attention performance (measured by conflict effects) as criterion variable for the male and female sex/gender group separately. In both models, we controlled for age because previous literature indicates detrimental age effects on executive functions (Ferguson et al., 2021).

**Regression model per sex/gender:**

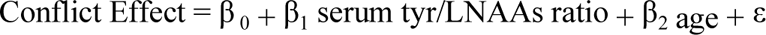

Further, a simple linear regression approach was applied to examine possible associations between age and dietary tyr intake, serum tyr levels or executive attention performance (Table 2). Regression analyses were performed including all participants. To test whether the effect of the variable age depended on the variable sex/gender, we extended the simple regression models by the interaction term age*sex/gender and repeated the analysis (**Fehler! Verweisquelle konnte nicht gefunden werden.**).

**Table 2:** Sensitivity Analyses Models: Dietary and Serum tyr

| **Models: Dietary - Serum tyr Relationship** |
| --- |
| Serum tyr Level = $\beta_0 + \beta_1$ Dietary tyr Intake + $\beta_2$ sex/gender + $\beta_3$ age + $\varepsilon$ |
| Serum tyr Level = $\beta_0 + \beta_1$ Dietary tyr Intake + $\beta_2$ sex/gender + $\beta_3$ age + $\beta_4$ BMI + $\varepsilon$ |
| Serum tyr Level = $\beta_0 + \beta_1$ Dietary Tyrosine Intake + $\beta_2$ sex/gender + $\beta_3$ age + $\beta_4$ BMI + $\beta_5$ SES + $\varepsilon$ |

To further elucidate the neural basis of age effects, we performed voxel-wise whole-brain analyses to correlate FA values with age.

**General Linear Model:**

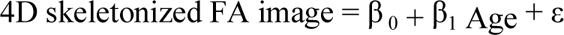

## Results

### Descriptives

In total, 59 individuals (female sex/gender group = 20, male sex/gender group = 39) with a mean age of 28.3 years (SD = 6.57), a mean BMI of 27.25 kg/m^2^ (SD = 1.48) and a mean SES score of 14.54 (SD = 3.11; n=51), reflecting medium to high socio- economic status, were included in the analyses (Table 3 & Figure 2).

**Figure 2:**
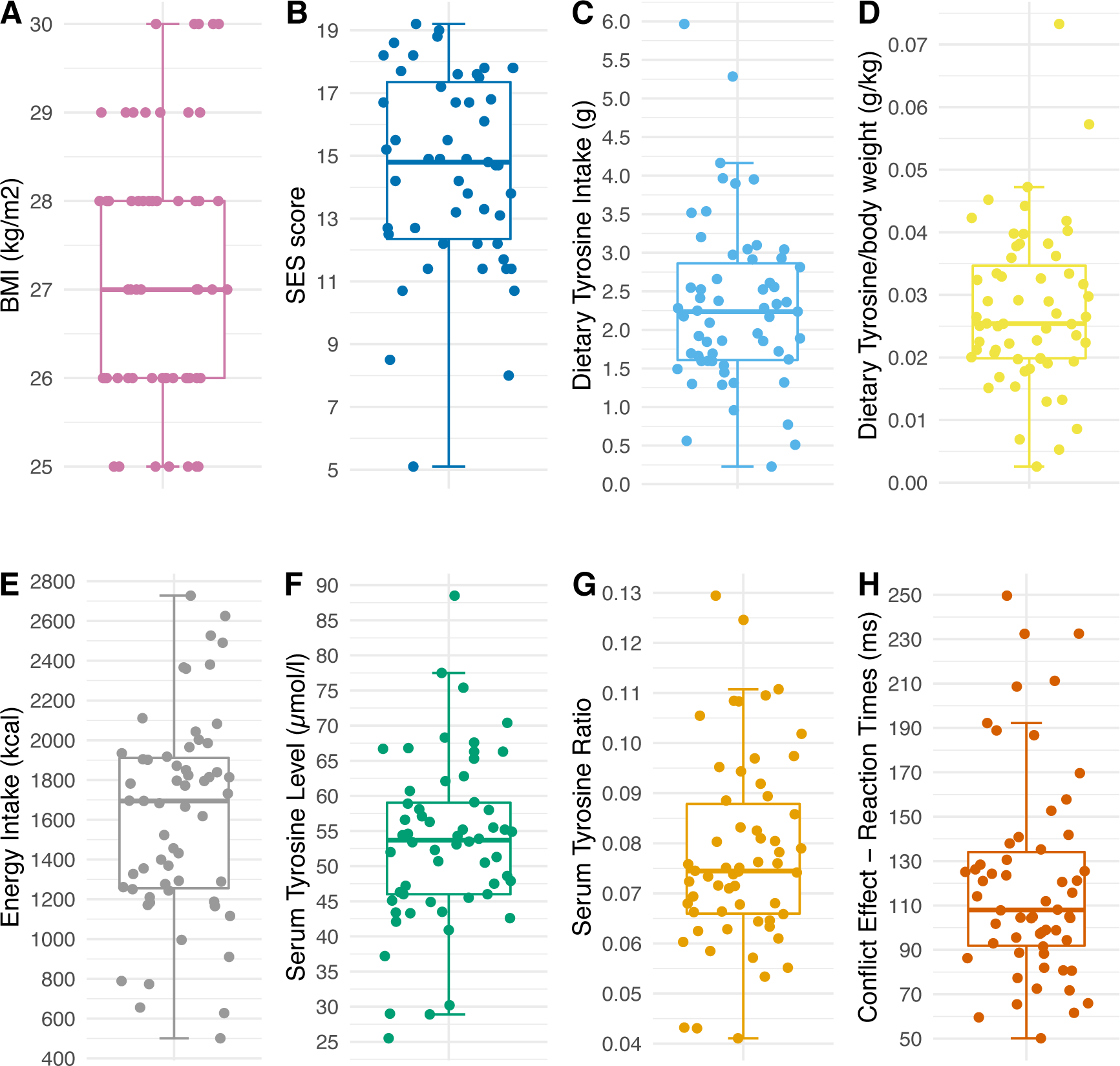
Descriptive statistics of variables of interest. Boxplots visualize maximum, minimum, median, first and third quartiles. The length of the whisker is 1.5 times the interquartile range (IQR); outliers are depicted above or below. Abbreviations: BMI, Body Mass Index; SES, Socio-Economic Status.

**Table 3:**
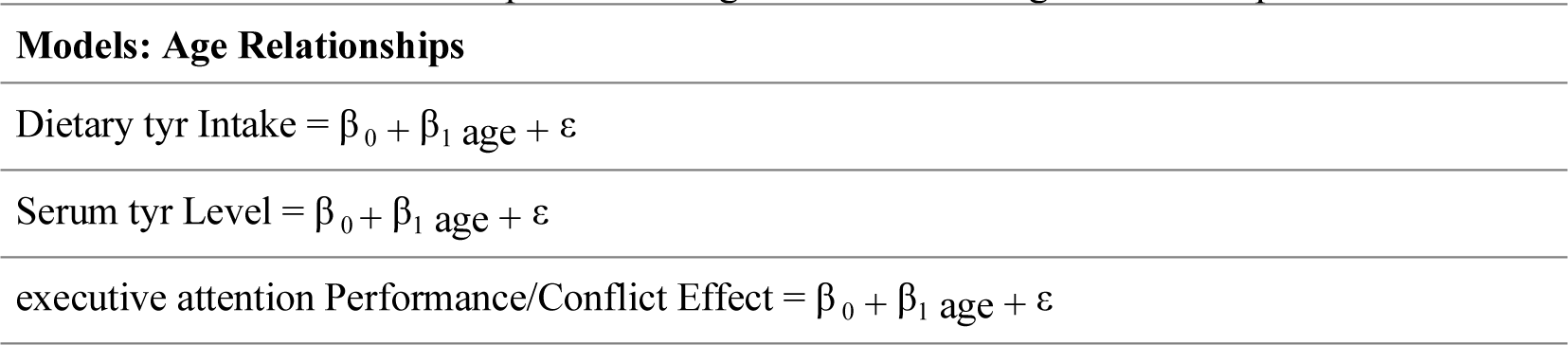
Simple Linear Regression Models: Age Relationship

On average participants consumed 2.3 g of tyr per day (SD = 1.07) or 0.03 g tyr per kg of body weight per day (SD = 0.01) and differed greatly in total energy intake (M =1615.9, SD = 509.51; min = 501 kcal/d; max = 2727 kcal/d). Serum tyr levels ranged from 25.5 to 88.5 µmol/l (M = 53.37, SD = 11.99) and serum tyr/LNAAs ratios (M = 0.08, SD = 0.02) from 0.04 to 0.13. Regarding executive attention performance quantified by conflict effects based on reaction times (RT), the sample showed a wide range from 50 ms to 250 ms with a mean conflict effect of 120.09 ms (SD = 45.49).

### ANT performance

The medium error rate was consistently 0% across all flanker and cue conditions with the exception of the incongruent flanker type (Figure 3 A & C). Both mean RT and mean error rate were highest in the incongruent flanker condition across all cue types (main effect of flanker type, error rate: F(2, 696) = 101.582, p < 2e^-16^, RT: F(2, 684) = 171.096, p < 2e^-16^; Figure 3 C & D) (see **SI** Table 8 **&** Table 9 for details). In addition, the spatial cue condition resulted in the lowest mean RT and the no-cue condition resulted in the highest mean RT across all flanker conditions (F(3, 684) = 26.241, p = 4.49e^-16^, Figure 3 D). Overall, these results reflected the pattern reported by the original study by Fan and colleagues (Fan et al., 2002).

**Figure 3:**
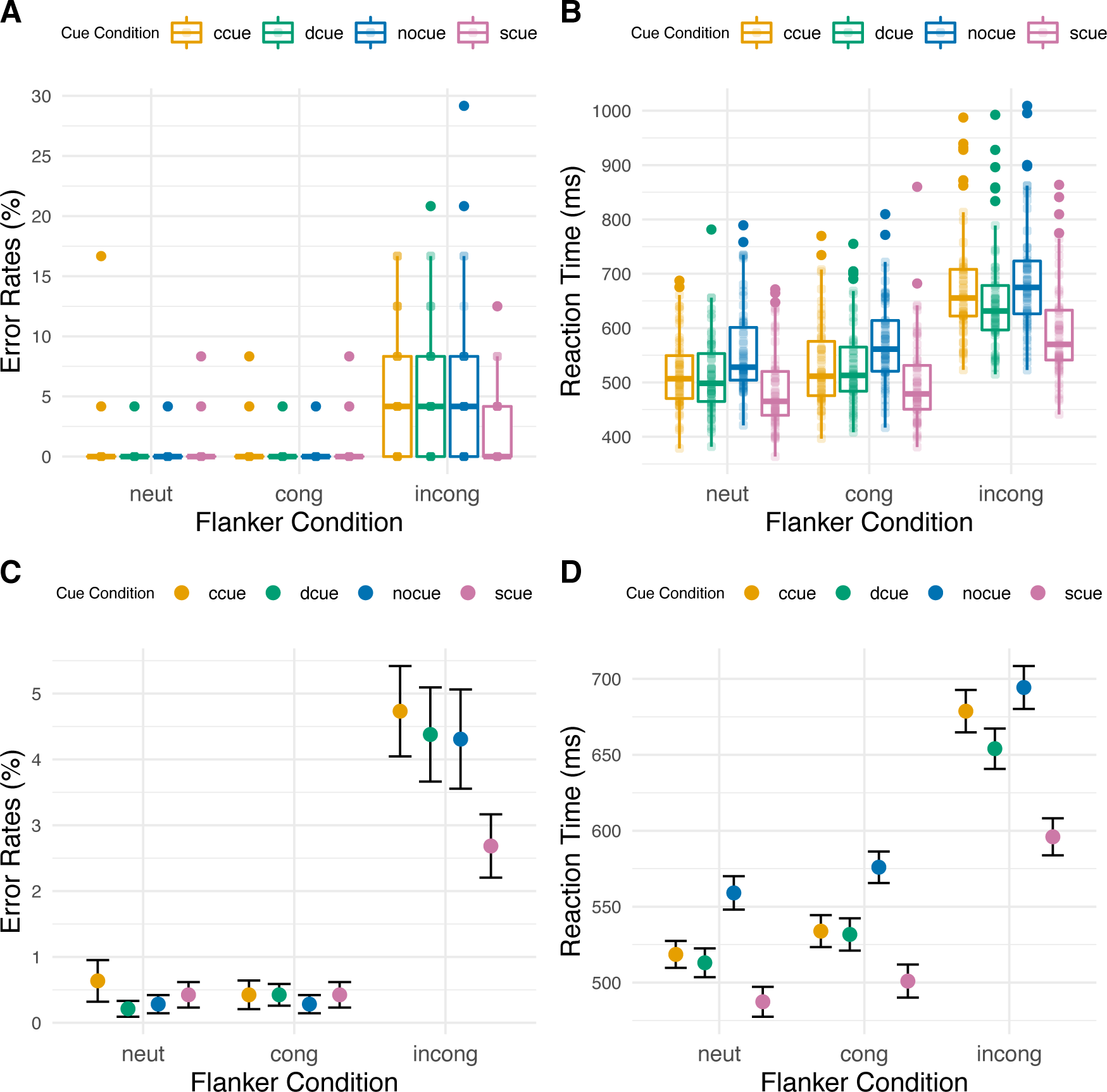
Error rates and reaction times as a function of cue condition and flanker type. Figure A & B visualize minimum, maximum, median, first and third quartiles. The length of the whiskers is 1.5 times the interquartile range (IQR); outliers are depicted above or below. Figure C & D show means and standard errors. Abbreviations: ccue, center cue; dcue, double cue; nocue, no cue; scue, spatial cue; neut, neutral; cong, congruent; incong, incongruent.

Post-hoc Tukey tests for RT showed that the difference in means is highly significant for the incongruent vs. neutral and incongruent vs. congruent flanker type pairs and for all cue-type pairs (p_adj_ < 0.003), except for the congruent vs. neutral flanker type pair (p = 0.11) and the double cue vs. center cue pair (p_adj_ = 0.65).

Summary statistics considering alerting, orienting and conflict (executive attention) effect of the ANT are summarized in Table 4 and visualized in Figure 4. Note that the mean conflict (executive attention) effect of 120 ms (SD = 46) was larger than that measured by Fan et al. (84 ms, SD = 25).

**Figure 4:**
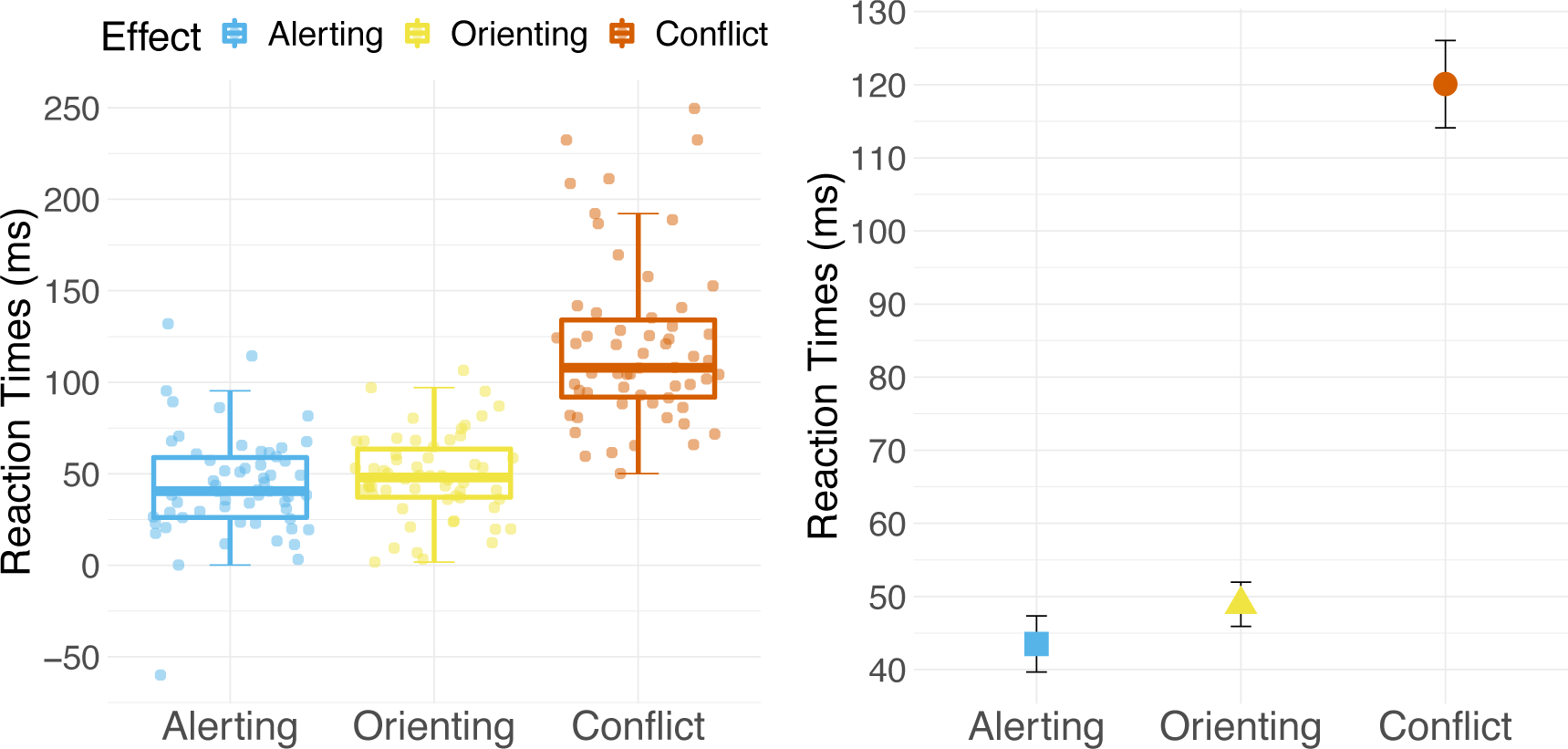
Summary statistics of ANT effects (i.e.; alerting effect, orienting effect, conflict effect) based on reaction time data. First figure visualizes minimum, maximum, median, first and third quartiles. The length of the whiskers is 1.5 times the interquartile range (IQR); outliers are depicted above or below. Second figure shows means and standard errors. Abbreviations: ANT, attention network test.

**Table 4:**
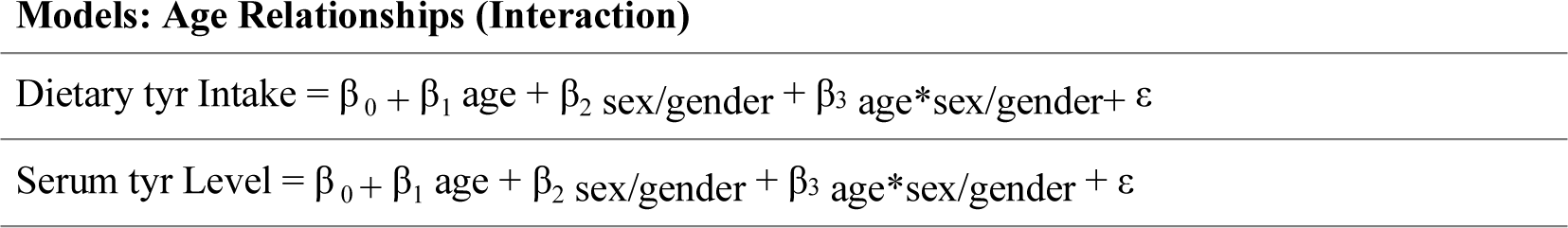
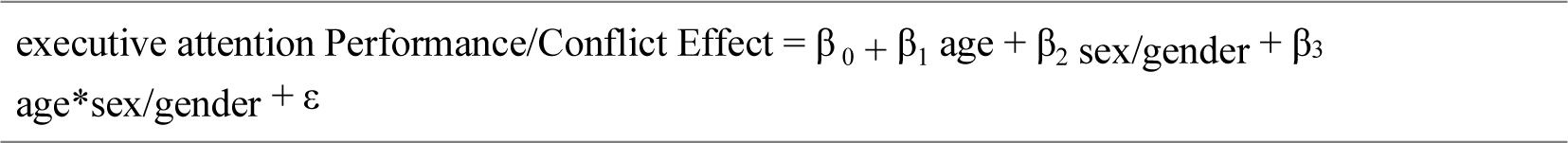
Multiple Regression Models: Age Relationships

### Dietary - serum tyr relationship (pre-registered)

In contrast to our hypothesis, dietary tyr intake was not significantly related to serum tyr levels according to linear regression (β = 0.198, p = 0.89). However, the model revealed sex/gender as a significant predictor (β = 13.07, p = 9.58e-05; R_adj_^2^ = 0.25, F(2, 55) = 10.31, p < 0.0002).

### Serum tyr ratio and executive attention performance (pre-registered)

Against our expectation, serum tyr ratios were not significantly related to executive attention performance (β = 0.42, p = 0.999). However, the non-significant model (Rad ^2^ = 0.05, F(2, 54) = 2.42, p < 0.098) pointed towards sex/gender as significant predictor of executive attention performance (β = -27.50, p < 0.04).

### Executive attention and white matter microstructure (pre-registered)

Two individuals were excluded due to missing ANT or dwMRI data. With respect to executive attention performance operationalized as conflict effects, results show neither a positive (TFCE, pFWE > 0.16) nor a negative (TFCE, pFWE > 0.95) relationship between differences in white matter integrity and executive attention performance.

### Sensitivity analyses (pre-registered)

First, a regression model with absolute serum tyr level as criterion and dietary tyr as predictor variable was still statistically significant when age was included in the model as an additional control variable (Rad ^2^ = 0.23, F(3, 54) = 6.78, p < 0.001). Neither dietary tyr nor age did predict absolute serum tyr level (age: β = 0.06, p < 0.81). However, the model still showed sex/gender as a significant predictor (β = 12.92, p < 0.0002).

This was similar when adding subsequently BMI and SES into the model (full model: R_adj_^2^ = 0.15, F(5, 44) = 2.67, p < 0.04). Neither age (β = 0.17, p = 0.57) nor BMI (β = 0.14, p < 0.91) nor SES (β = - 0.12, p < 0.83) significantly predicted absolute serum tyr levels. However, the variable sex/gender remained a significant predictor (β = 12.08, p < 0.002).

When including age in the regression model to control for general effects of aging on executive attention performance, the overall model remained non-significant (Rad ^2^ = 0.06, F(3, 53) = 2.24, p < 0.095) and age did not significantly predict executive attention performance (β = 1.25, p = 0.19). Again, the non-significant model favoured sex/gender as a predictor (β = -32.49, p < 0.02). This was similar when subsequently adding BMI and SES into the model, in the full model, neither age (β = 0.12850, p < 0.9046) nor BMI (β = -0.13958, p < 0.9751) nor SES (β = 0.046, p < 0.98) were significantly related to executive attention performance, but sex/gender was (β = -29.38, p < 0.03).

We also used linear regression to examine whether daily dietary tyr intake (adjusted for body weight) was linked to executive attention performance when controlling for total energy intake. Again, the overall regression model did not indicate a link between dietary tyr intake adjusted for body weight and executive attention performance (R_adj_^2^ = -0.02325, F(2, 55) = 0.3524, p = 0.7046), indicating that differences in executive attention performance could not be explained by differences in energy intake adjusted dietary tyr intake (β = -0.047, p < 0.77).

### Exploratory analyses

Dietary tyr intake did not correlate with either absolute tyr levels (R = 0.2, p = 0.14) or relative serum tyr/LNAAs ratios (R = 0.19, p = 0.16), nor when tyr intake was adjusted for energy intake (R = 0.18, p = 0.19) or body weight (R = 0.0098, p = 0.46) (Figure 5). However, higher absolute serum tyr levels related to higher relative serum tyr ratios (R = 0.52, p =2.6 e^-05^) (Figure 6).

**Figure 5:**
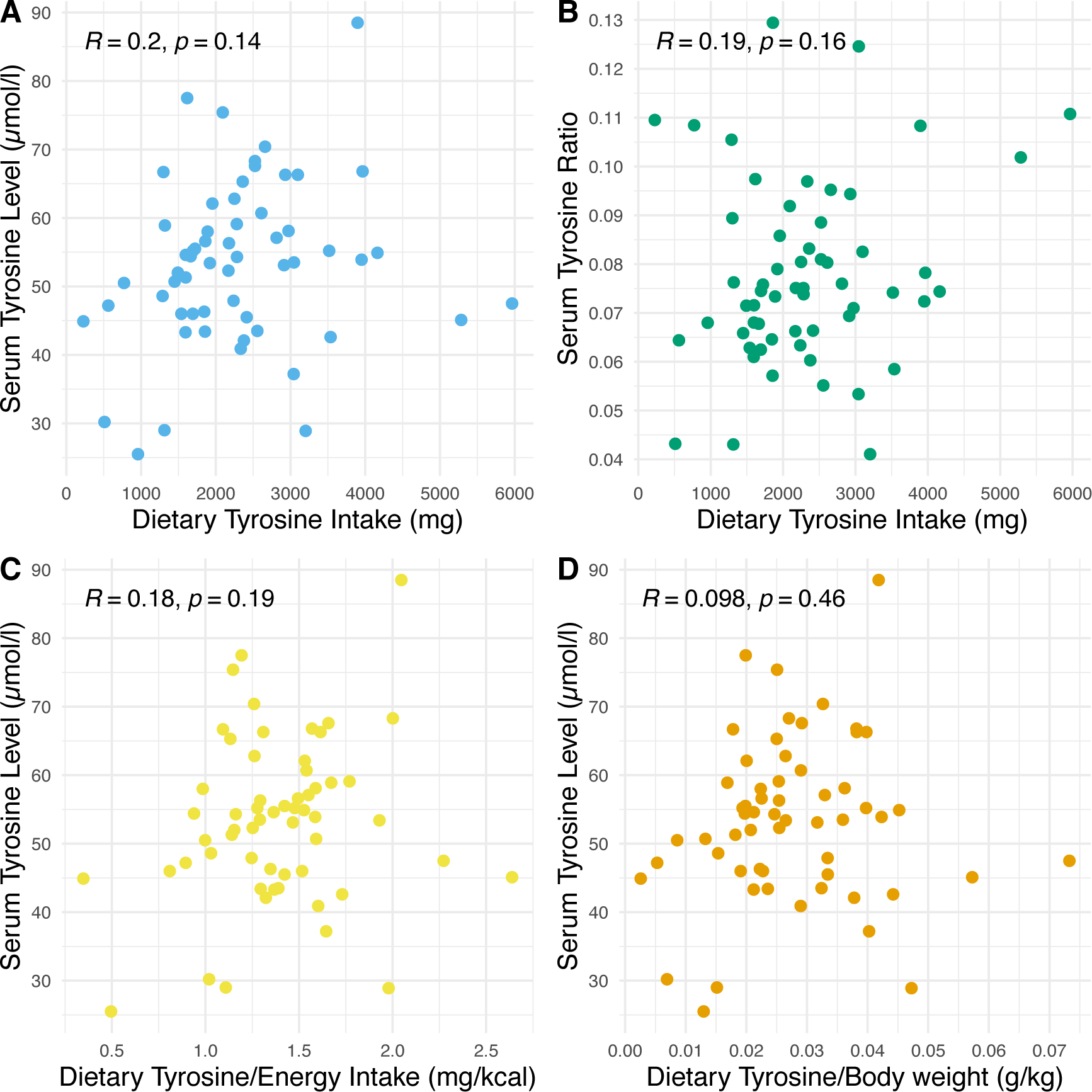
Correlations of (A) dietary tyrosine intake and absolute serum tyrosine levels, (B) dietary tyrosine intake and relative serum tyrosine/LNAA ratios, (C) dietary tyrosine intake adjusted for energy intake and absolute serum tyrosine levels, (D) dietary tyrosine intake adjusted for body weight and absolute serum tyrosine levels. Abbreviations: LNAA; large neutral amino acids; R, Pearson’s Correlation Coefficients; p, p-value.

**Figure 6:**
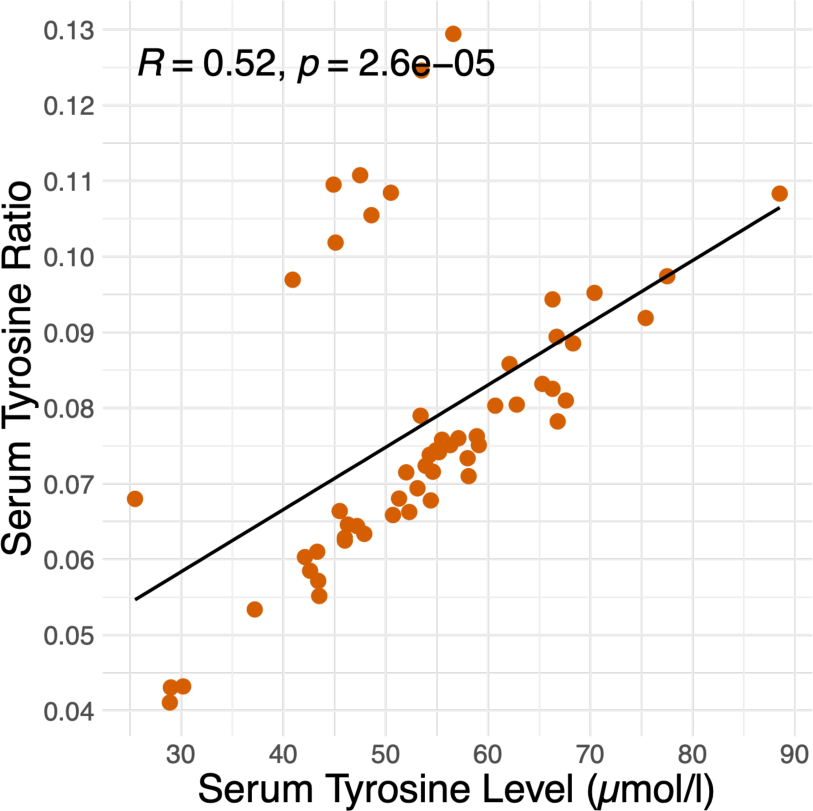
Correlation of absolute serum tyrosine levels and relative serum tyrosine/LNAA ratios. Abbreviations: LNAA; large neutral amino acids; R, Pearson’s Correlation Coefficients; p, p-value

Further, we observed significant sex/gender difference in dietary tyr intake, with men having higher values than women (W = 236, p = 0.02) (Figure 7). However, the difference was no longer significant when dietary tyr intake was adjusted for total energy intake (W = 285, p = 0.09), as total energy intake was also significantly different between sex/gender groups (W = 252, p = 0.03) (Figure 7).

**Figure 7:**
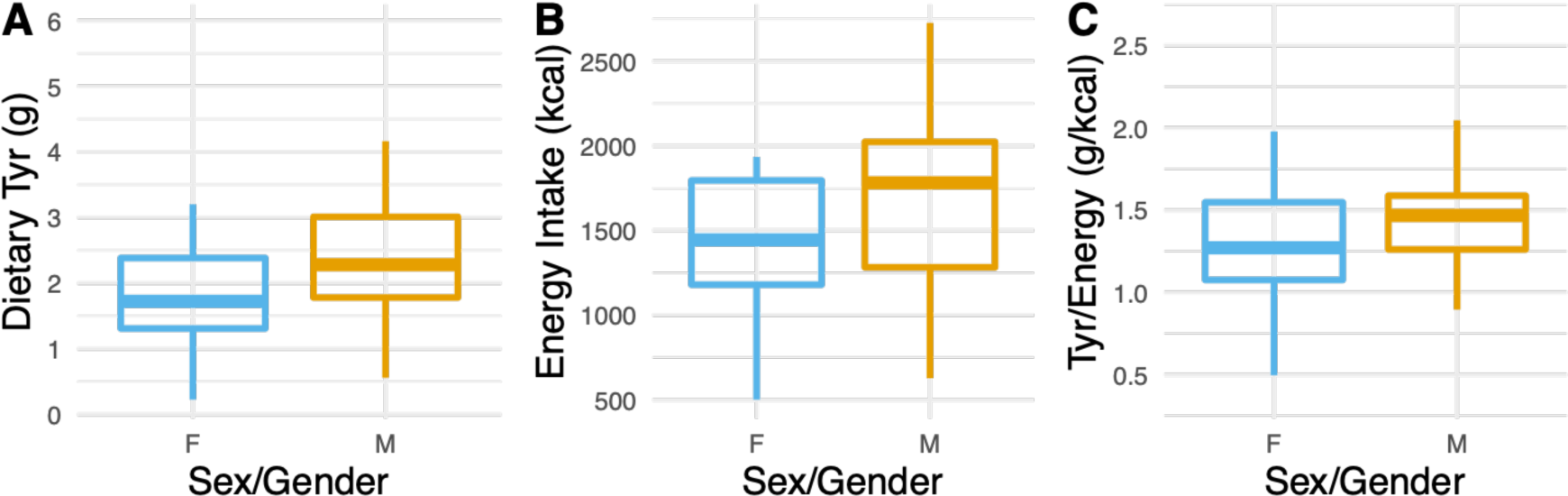
Sex/gender group differences regarding the variables of interest; Figure A shows the significant sex/gender group difference in dietary tyrosine intake; Figure B and C depict non-significant sex/gender group differences in energy intake and in dietary tyrosine intake adjusted for total energy intake. Abbreviations: Tyr, Tyrosine; TYR/Energy, Dietary Tyrosine Intake adjusted for total energy intake.

Furthermore, the male sex/gender group showed better executive attention performance operationalized as conflict effects based on reaction times (W = 493, p = 0.04), although the female sex/gender group was significantly younger than the male sex/gender group (W = 211, p = 0.004). Moreover, our results showed that the male sex/gender group had significantly higher serum tyr markers than the female sex/gender group, both in terms of absolute tyr levels (W = 118, p < 0.00003) and serum tyr/LNAAs ratios (W = 232, p = 0.02) (Figure 8). The two sex/gender groups did not differ significantly in either SES scores (W = 274, p = 0.76) or BMI (W = 331, p = 0.35).

**Figure 8:**
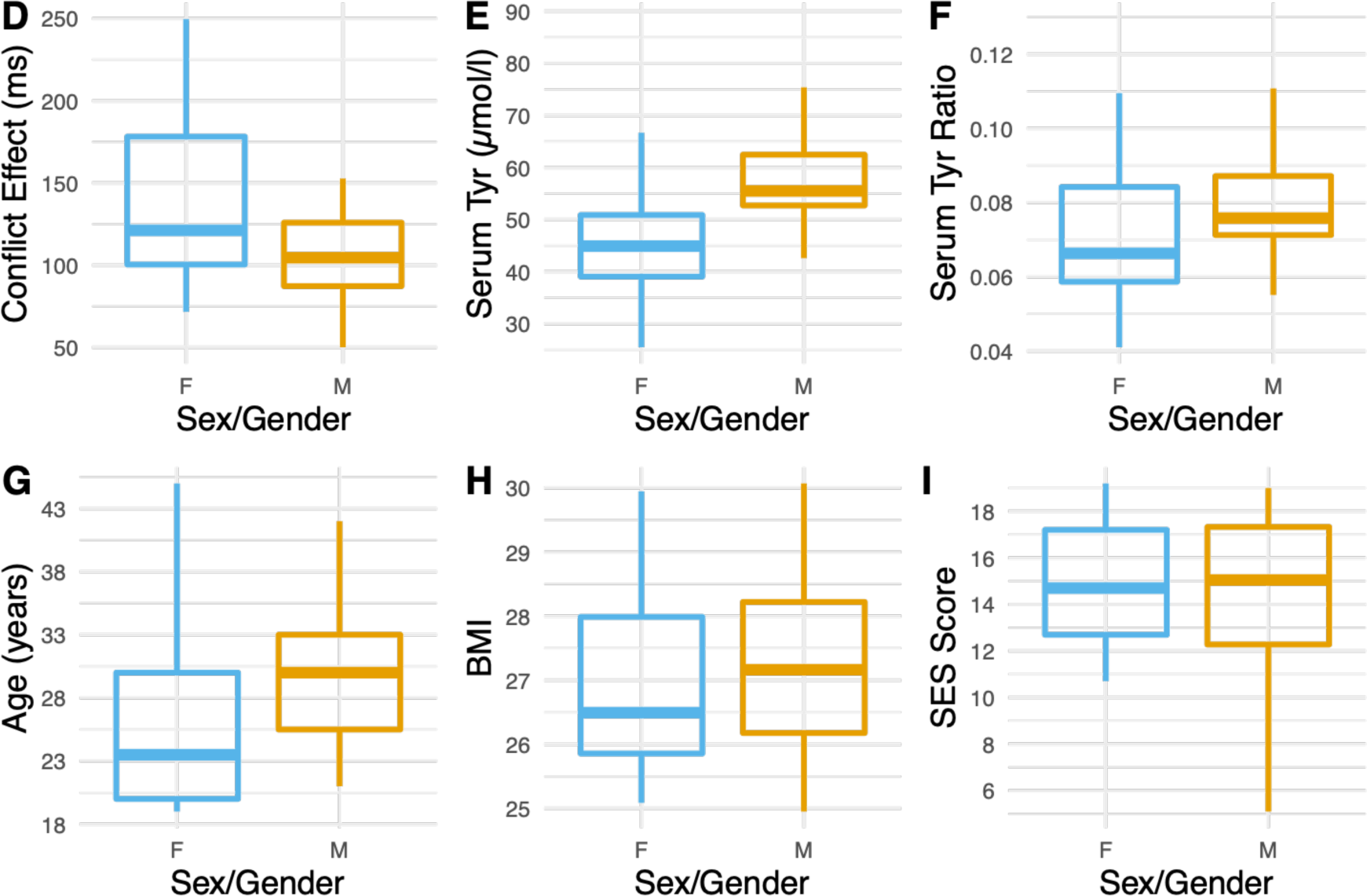
Significant sex/gender group differences in executive attention performance (i.e., conflict effect) (Figure D), absolute serum tyrosine levels (Figure E), relative serum tyrosine ratios (Figure F) and age (Figure G). Non-significant sex/gender group differences in BMI (Figure H) and SES Score (Figure I). Abbreviations: Tyr, Tyrosine; BMI, Body Mass Index; SES, Socio-Economic Status. Boxplots depict median as line and Q1 and Q3 as boxes.

In the female sex/gender group, we observed that neither serum tyr/LNAAS ratios nor age were related to conflict effects (R_adj_^2^ = 0.01, F(2, 15) = 1.098, p < 0.36). In the male sex/gender group, however, serum tyr ratio significantly predicted conflict effects (β = - 906.95, p < 0.01), when taking the effect of age into account (β = 3.39, p < 0.001).

According to the model, conflict effects deteriorated by a RT of 3.39 ms with each additional year of life (Table 5). In contrast, the positive influence of tyr in terms of an improvement of executive attention performance by more than 900 ms per increase of 1 in the ratio appeared surprisingly large. However, a serum tyr/LNAAs ratio of 2 means that twice as much tyr must be present in the blood relative to the sum of all other LNAAs. Therefore, this number represents rather a theoretical value.

**Table 5:**
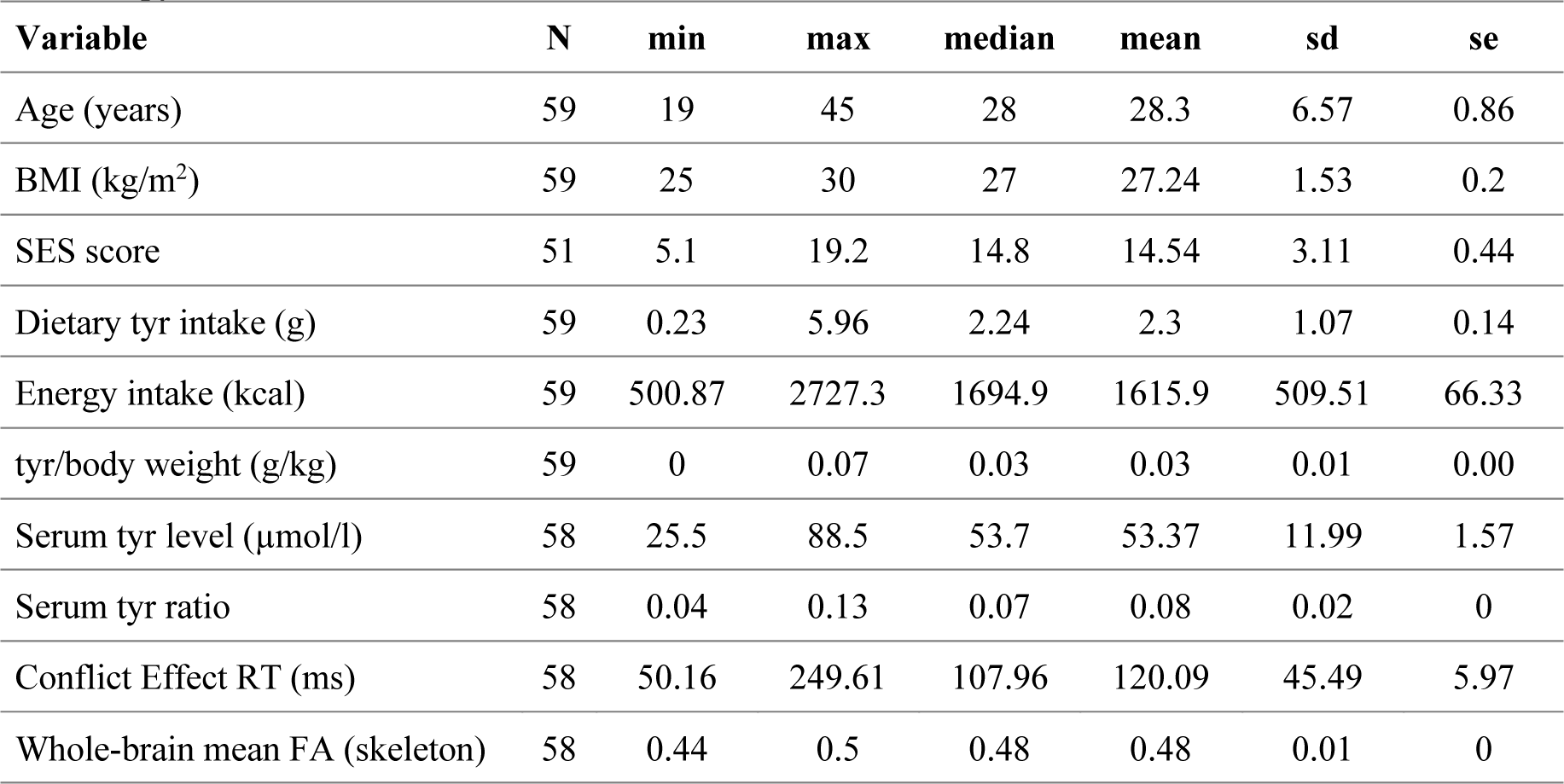
Descriptive Statistics of the sample. Abbreviations: n, number of subjects; BMI, Body Mass Index; SD, standard deviation; SE, standard error; SES, socio-economic status; RT, reaction time; FA, fractional anisotropy.

Higher age was associated with higher dietary tyr intake (R_adj_^2^ = 0.1, F(1, 57) = 7.57, p = 0.0079) in our sample (Figure 9), and the effect of age on executive attention performance was dependent on sex/gender (R_adj_^2^ = 0.16, F (3, 54) = 4.48, p = 0.007) (Figure 10 & Table 6).

**Figure 9:**
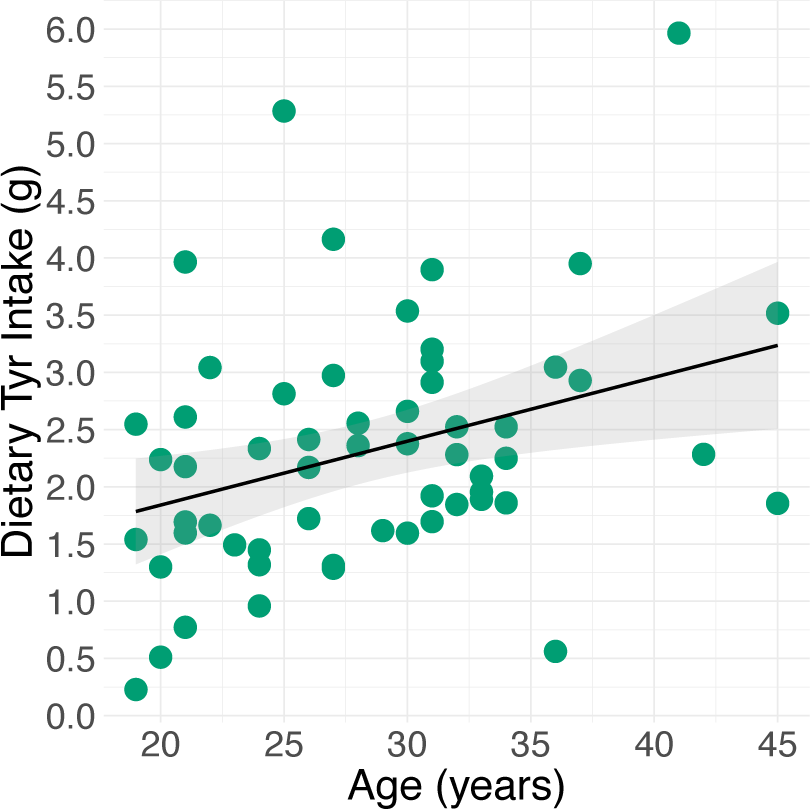
Age - Dietary Tyrosine Intake Relationship. The regression line is indicated in black. Abbreviations: Tyr, Tyrosine.

**Figure 10:**
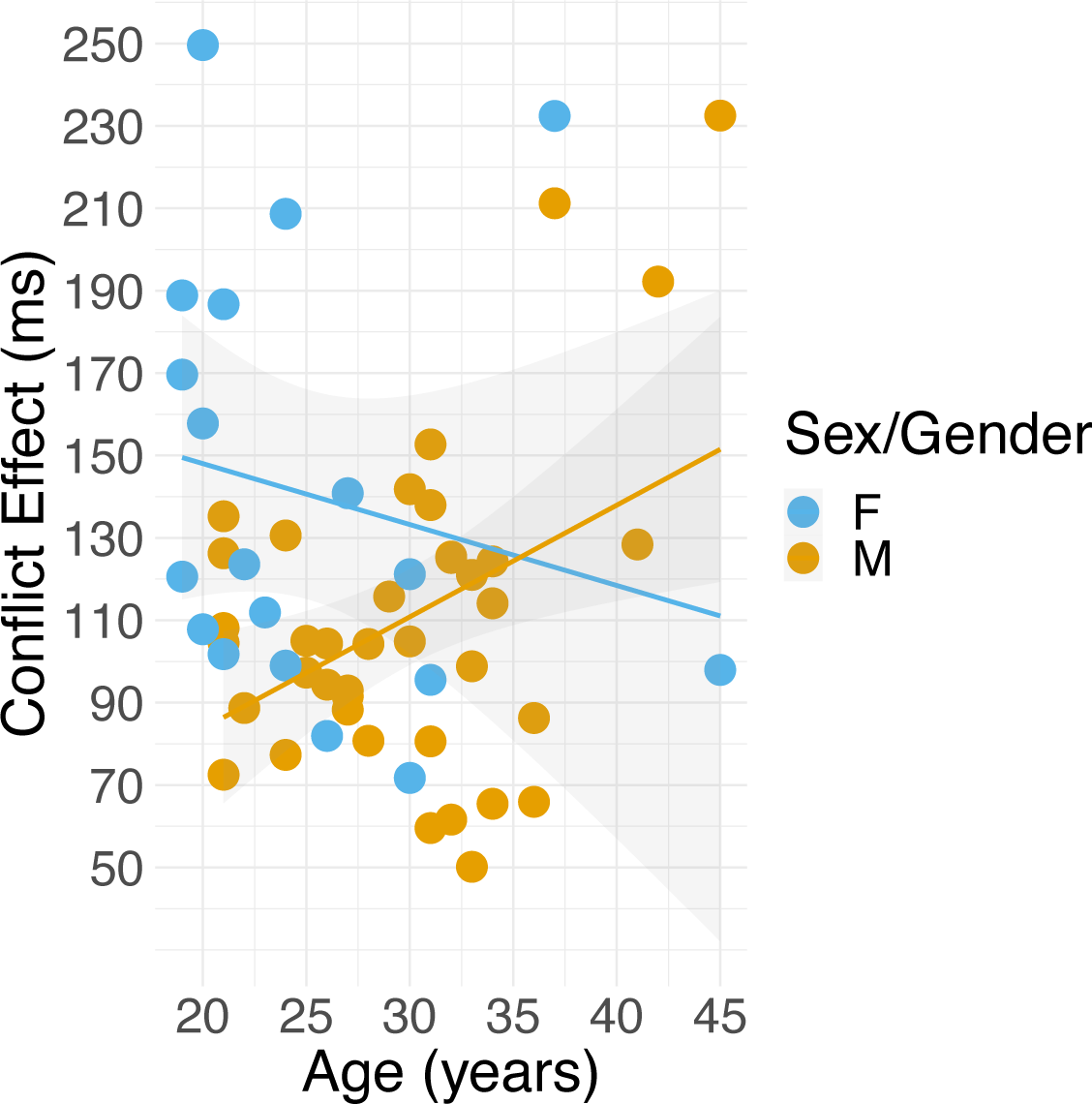
Moderation of the association between age and executive attention performance by sex/gender group. Abbreviations: F, female sex/gender group; M, male sex/gender group).

**Table 6:**
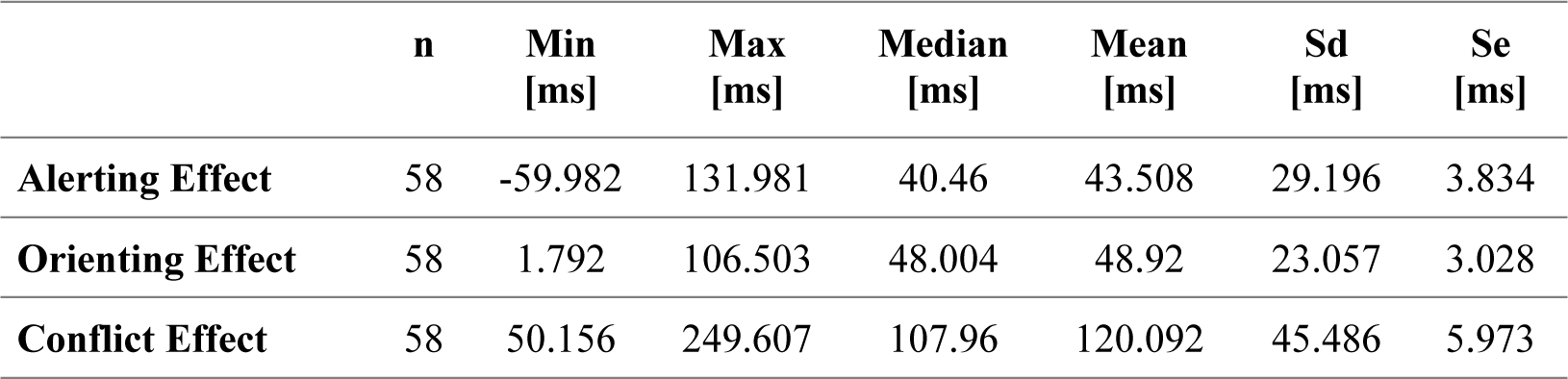
Summary Statistics ANT Effects. Abbreviations: n, number of subjects; sd, standard deviation; se, standard error.

To examine age-structure correlations, we found that higher age related to lower FA values (TFCE, pFWE > 0.0018), Figure 11), in seven significant clusters (Table 7), with the seventh being the largest cluster (number of voxels = 35226) widely distributed across the brain (Figure 12).

**Figure 11:**
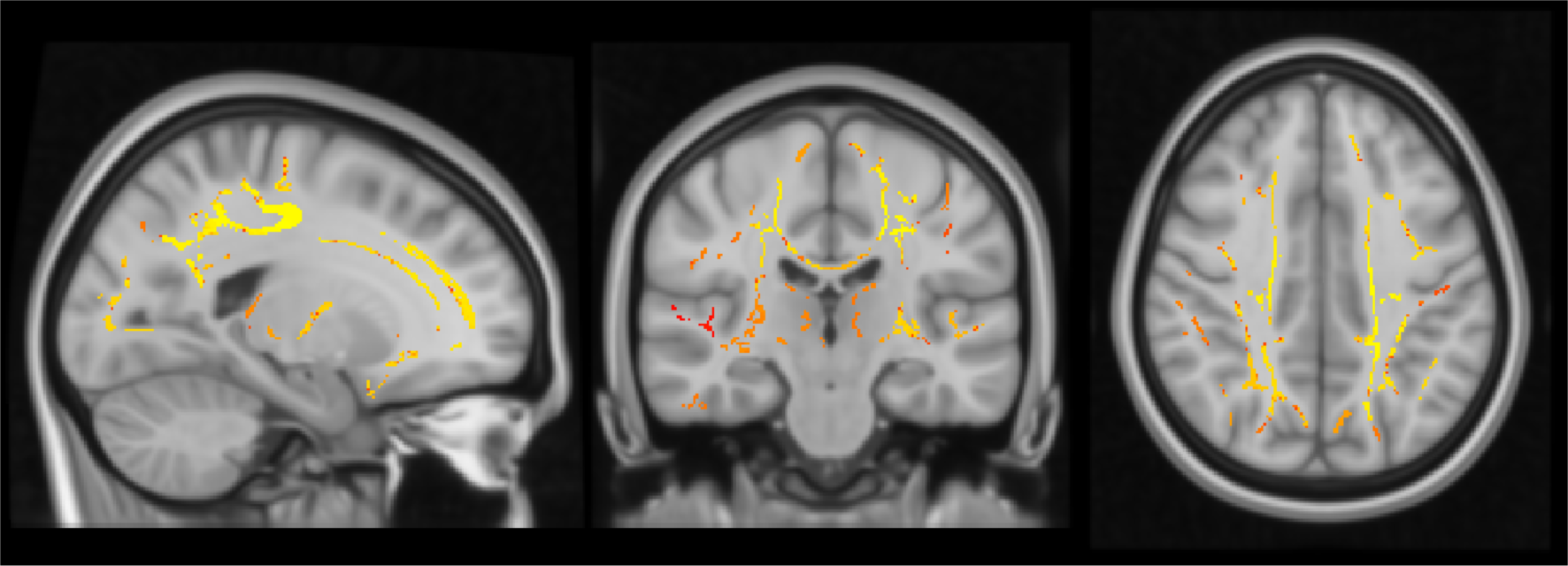
Whole-brain age-FA correlations (max(pFWE) = 0.05 (red), min(pFWE) = 0.0018 (yellow)) overlaid on the MNI152 template (coordinates of max. voxel. x = -20, y = -22, z = 39). Abbreviations: FA, fractional anisotropy; FWE, family-wise error corrected.

**Figure 12:**
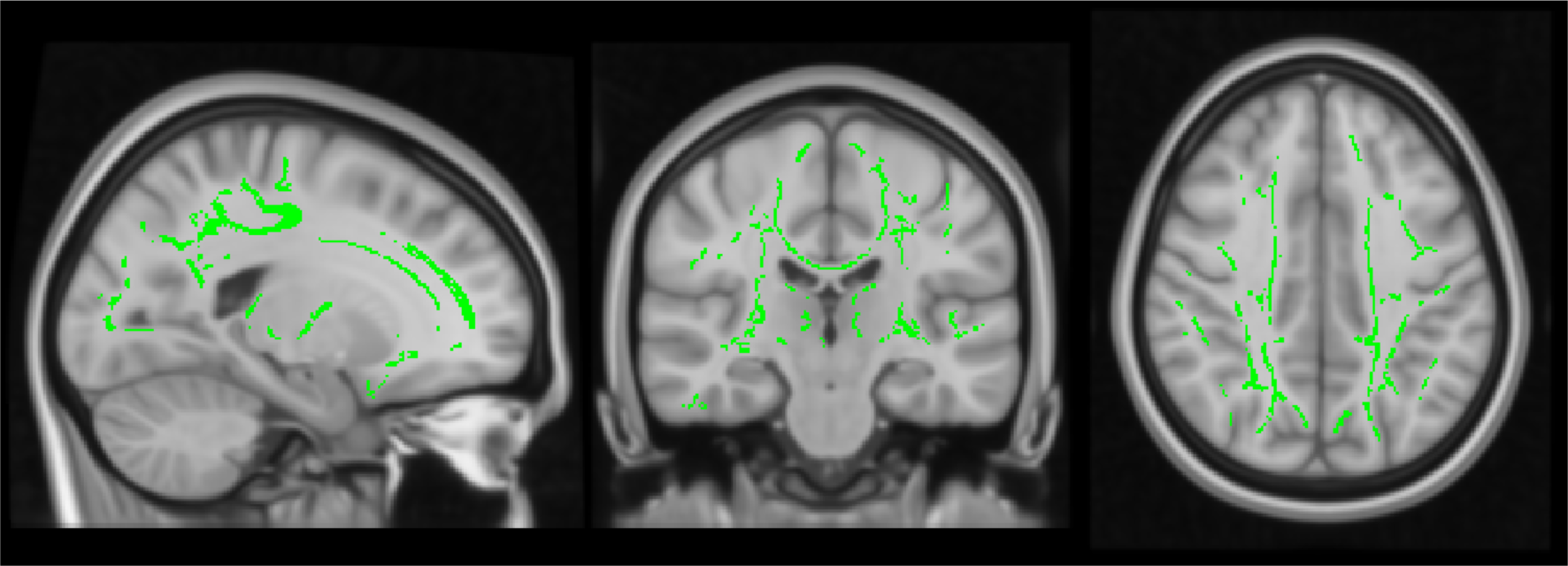
Maximal age-associated cluster (pFWE < 0.05; number of voxels = 35226) overlaid on the MNI152 template (coordinates of max. voxel. x = -20, y = -22, z = 39). Abbreviations: FWE, family-wise error corrected.

**Table 7:**
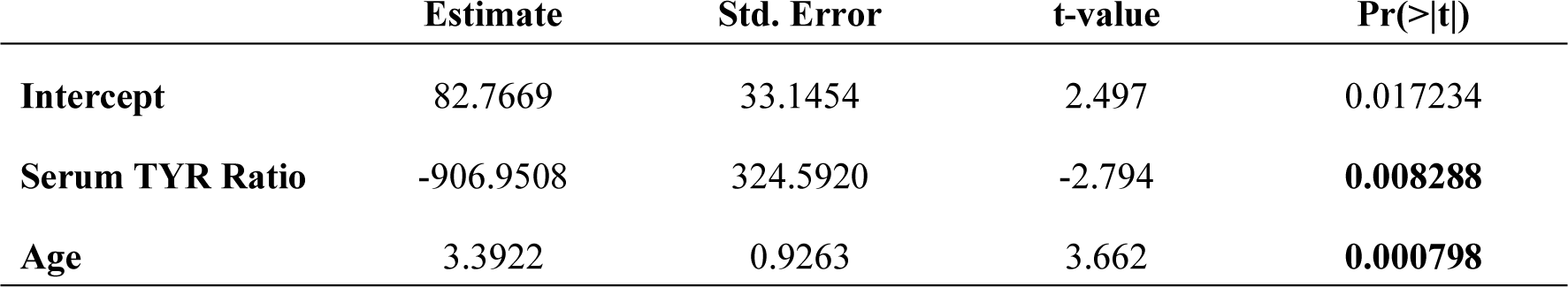
Coefficients of the Serum Tyrosine Ratio – Executive Attention Performance Regression Model controlling for age (only male sex/gender group). Abbreviations: Tyr, Tyrosine; Std. Error, Standard Error.

**Table 8:**
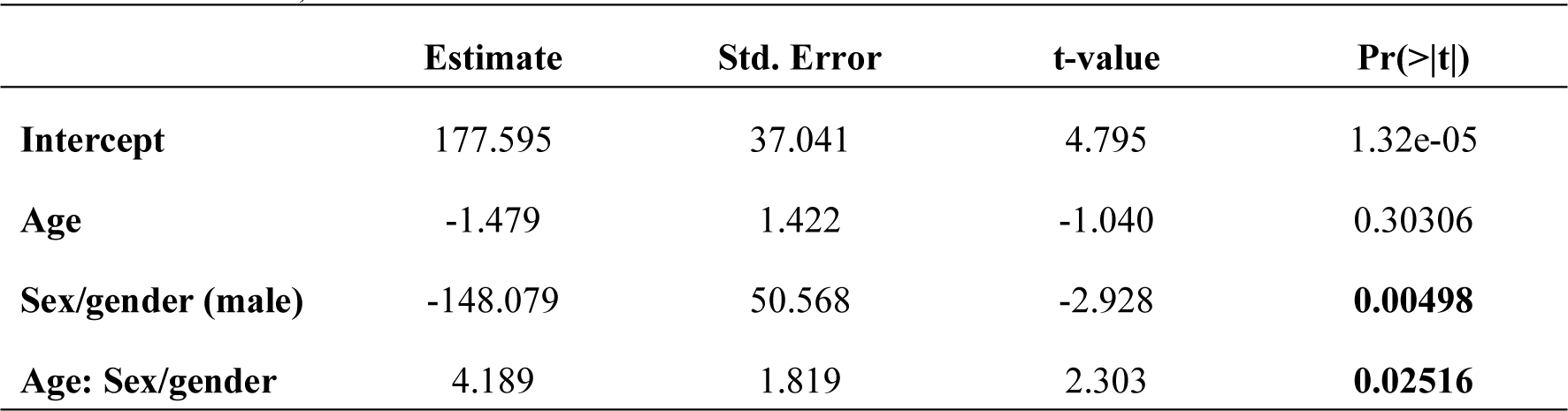
Coefficients of the Age - Executive Attention Performance Regression Model (with age:sex/gender interaction). Abbreviations: Std. Error, Standard Error.

## Discussion

In this cross-sectional analysis in 59 healthy, overweight adults, our pre-registered confirmatory analyses did not support any of the four hypotheses raised. Neither a relationship between dietary tyr intake and serum tyr levels nor between serum tyr/LNAAs ratios and executive attention performance could be demonstrated. In addition, whole-brain correlation analysis revealed no structure-function associations between white matter microstructure and executive attention performance. Therefore, no further moderation analyses could be performed. Finally, the results of the pre-registered exploratory analysis also did not indicate an association between self-reported dietary tyr intake and cognitive performance in our sample. Further exploratory analysis indicated that dietary and serum tyr as well as conflict effects were higher in the male sex/gender group and that older age was associated with higher dietary tyr intake and lower fractional anisotropy in a widespread cluster across the brain. Finally, a higher relative serum tyr/LNAA concentration was linked to better executive attention performance in the male sex/gender group when negative age effects were taken into account.

### Dietary and serum tyr

The measured dietary and serum tyr levels resembled those already described in studies with much larger sample sizes. For example, Kühn et al. reported a mean dietary tyr intake of 2.85 g per day, irrespective of the lower mean BMI (M =23.39 kg/m^2^) of the sample (341 younger adults, 53% women, mean age: 31, recruited from the Berlin Aging Study II (Kühn et al., 2019). Furthermore, mean serum tyr level of 53.37 µmol/l in our sample is close to the value obtained in a recently published large epidemiological study (M= 597 µmol/l, SD = 16.7, 1324 adults, 87% women, mean age: 44, recruited from the Australian Children’s B Cohort) in a sample with similar median BMI (26.54 kg/m^2^) (Andraos et al., 2021). The similarity of our sample to larger studies supports the reliability of our measurements and the methodology we used. Moreover, the result patterns of the ANT are largely consistent with those reported by the original study (Fan et al., 2002) indicating that the ANT has been properly performed and analyzed. However, analysis of the error rate data showed a ceiling effect. This might be insightful with regard to our healthy, young to middle-aged sample with medium to high socioeconomic status and presumably higher education.

With respect to the methods, self-reported data have been severely criticized as reliable measure of dietary behavior; especially with respect to energy underreporting due to recall biases (Subar et al., 2015). Yet, by carefully evaluating previous research findings and urging researchers to interpret results appropriately, consider limitations, and to improve methods, self-reported dietary measures seem valid. Moreover, the correlation of self-reported dietary intake with nutritional biomarkers has also been problematized (Naska et al., 2017). In particular, short-term concentration biomarkers (e.g.; serum concentrations) are affected by individual differences in metabolism and socio-demographical, behavioral or health-related characteristics leading to inconsistent findings in diet-related research. Thus, Naska and colleagues suggest to validate the applied nutrition questionnaires by employing multiple methods simultaneously and incorporating new technologies. In conclusion, we will consider these recommendations in future studies of self-reported dietary measures to overcome the limitations addressed.

Sample size was not powered to the current cross-sectional analysis, and might have been too low to detect the hypothesized effects. We propose larger samples, data pooling and a priori power analyses to circumvent underpowered analyses.

### Executive attention and white matter microstructure

We could not confirm the findings of two cross-sectional studies examining the association between self-reported dietary tyr intake and cognitive performance (Hensel et al., 2019; Kühn et al., 2019). Both studies measured working memory and were based on large sample sizes (n > 280). In contrast, our pre-registered exploratory regression analysis did not demonstrate a relationship between dietary tyr intake and executive attention possibly due to a lack of power—the power calculation was based on effects regarding the intervention study; or possible differences in DA processing between executive attention and working memory.

Considering the null findings of the whole-brain voxel-wise correlation analysis, it could be argued that DTI analyses are more sensitive to group analyses and comparisons of patient vs. non-patient groups rather than at the individual level in correlations analysis (Op de Beeck & Nakatani, 2019). Moreover, we decided against a region of interest (ROI) approach and in favor of a whole-brain analysis to explore structure-function relationships without local restriction, although previous literature suggested the anterior corona radiata (ACR) as interesting ROI regrading executive attention performance (Niogi et al., 2010; Yin et al., 2013) and thus potentially suffered from lower statistical power. In further analyses, we aim at employing a ROI approach to increase statistical power by decreasing corrections for multiple comparisons (Saxe et al., 2006) by defining ROIs in advance, fewer statistical tests need to be performed and subsequently fewer corrections for multiple comparisons need to be made.

### Exploratory findings

Considering exploratory analyses, sex/gender emerged as significant predictor of serum tyr concentrations. This unexpected finding also suggests that other relevant aspects influencing the variables of interest should be included in the analyses elucidating the link between diet and cognition. Furthermore, exploratory analyses showed sex/gender-specific differences with respect to various variables of interest: an effect of age on dietary tyr intake and sex/gender-age interaction effects in relation to executive attention performance. In addition, whole-brain correlation analysis showed that older age was associated with lower FA in a widespread cluster. Interestingly, we demonstrated a significant association between relative tyr serum levels and executive attention performance in the male sex/gender group only when taking negative age effects on white matter into account.

Correlation analysis of various dietary and serum tyr markers showed no significant associations. Thus, the results call a direct relationship between self-reported subjective measures of dietary tyr intake and objectively measured serum tyr levels into question. However, based on these null results, tools like FFQ based on self-report should not be doubted as a reliable method for measuring dietary tyr intake, but should rather be used as an opportunity to implement recommendations to improve future studies of self-reported dietary measures. In contrast to the non-significant diet-serum associations, we demonstrated a moderate to strong correlation between the two serum markers, suggesting that individuals with higher tyr levels also had higher levels of all other six LNAAs.

### Considering sex/gender differences

We demonstrated various sex/gender differences and sex/gender-age interaction effects. Although numerous research findings point to a wide variety of sex/gender differences in brain anatomy and function (Bale, 2019; Ruigrok et al., 2014; Spets & Slotnick, 2020), the investigation and reporting of sex/gender differences is a topic of ongoing debate in medical and neuroscientific research. Biological sex differences are commonly neglected in clinical research, resulting in significant disadvantages for patients with female biological characteristics due to incorrect dosing of medications or unanticipated side effects (Cahill, 2014). Not only is the biological male the default model in neuroscience research, but also fewer participants exhibiting female biological characteristics are being included in research studies to reduce complexity. Widespread misconceptions contribute to an underestimated relevance of investigating sex/gender differences in neuroscience (Cahill, 2006): sex/gender-specific influences are said to be merely small, unreliable and result from only a few extreme cases. However, the main concerns against research on sex/gender differences refer to implicit reinforcement of gender stereotypes and sexism by claiming “hardwired brain sex differences” (Jordan-Young & Rumiati, 2012; Persson & Pownall, 2021). Additionally, a potential reporting bias towards sex/gender differences may be present (Cahill, 2020; David et al., 2018). Most importantly, critics emphasize that the concepts of biological sex and social gender would be inextricably linked, making it a fluid, multi-layered, complex structure that might be closely interwoven with cultural and social factors and could not be reduced to biological and dichotomous differences (Jordan-Young & Rumiati, 2012).

We consider these concerns valid, not only given the problematic acquisition of the study variable sex/gender in the context of our own study design, but also with respect to various publications that use inaccurate definitions, operationalizations, and thus fuzzy interpretations of the obtained findings. Moreover, we also regard the classic model of sexual differentiation as outdated which should be replaced by the parallel model of sex/gender differentiation, which stresses the parallel action of genes, hormones, and the environment. Biological and sociocultural feminine or masculine aspects form a “mosaic” from which gender identity as a whole emerges (Rouse & Hamilton, 2021). Nevertheless, differences in biological or non-biological characteristics should not be fully negated. Since recently a paradigm shift is taking place, which is reflected in the approval of research funds on the topic of sex/gender and in a change in the editorial policy of relevant journals (Zsido & Sacher, 2021).

### Sex/gender differences in tyr - cognition relation

In line with our findings, several studies have reported sex/gender differences in eating behavior, food choices, and dietary strategies before (Arganini et al., 2012; Grzymisławska et al., 2020). Food intake for males and females differed not only in terms of quantity and quality, but also in terms of frequency, timing, and location of meals. For example, women are more likely to eat foods with higher fiber content (e.g., fruits and vegetables), while men tend to consume more dietary supplements. A reciprocal model of the relationship between sex/gender and diet has been proposed, shaped by the interplay of physiological, psychological, and sociocultural factors (Grzymisławska et al., 2020). Both dietary behavior and sex/gender are influenced by psychological factors (e.g., stress, mood), geophysical factors (e.g., local topography, climatic conditions, and infrastructure), socioeconomic factors (e.g., education, occupation, income, and marital status), and cultural factors (e.g., traditions, cultural heritage, and religion). Moreover, the variable sex/gender is not sufficiently considered in nutritional biomarker-related research (Song et al., 2018). Only 68% of the studies included in a review investigating the consideration of sex/gender in nutrition research determined nutrition-related biomarker indices by sex/gender. Of these, only 13% reported nutrition-related biomarker indices for men and women separately, 33% adjusted for sex/gender or performed analyses considering sex/gender without reporting sex/gender-specific results, and more than half (54%) did not consider the variable sex/gender at all. A recent review emphasizes the need to consider sex/gender differences in nutrition-related research because of sex/gender differences in serum metabolite concentrations and AA level dynamics during the menstrual cycle (Brennan & Gibbons, 2020). An epidemiological study reporting higher plasma tyr levels in men points to research indicating higher insulin concentrations and insulin sensitivity in women (Andraos et al., 2021), which is related to sex/gender differences in the amount and distribution of adipose tissue (Valencak et al., 2017). Insulin promotes the uptake of AA in peripheral tissues and thus decreases AA concentrations in blood and uptake at the blood-brain barrier (John D. Fernstrom, 2013). Strang and colleagues justified the selection of exclusively male subjects with sex/gender differences in metabolism (Strang et al., 2017) based on differences in abdominal adipose tissue. Sex/gender differences may also explain some of the previously published null results, e.g. no intervention effect of tyr supplementation on executive attention in females only (Frings et al., 2020).

Our results show higher serum tyr for the male sex/gender group compared to the female sex/gender group, which is consistent with the literature (Andraos et al., 2021; Darst et al., 2019). The male sex/gender group showed better executive attention performance, but only in terms of significantly lower conflict effects based on reaction times. However, accuracy scores did not significantly differ between the two sex/gender groups. This pattern is also consistent with the current literature on sex/gender differences in cognition. A literature review concluded that there is little to no evidence for a consistent difference in attention, impulsive action, decision making, and working memory (Grissom & Reyes, 2019). Therein, differences in slower reaction times and a tendency of females to avoid negative consequences in decision making were reported. In addition, men and women seemed to differ in their working memory strategies and underlying neural activity patterns, possibly due to differences in neurotransmitter systems and structural brain development. Another meta-analysis demonstrated minor sex/gender differences in executive control only at the individual task level, but not at the domain level (i.e., performance monitoring, response inhibition, and cognitive set-shifting) (Gaillard et al., 2021). In sum, all herein demonstrated sex/gender differences are not surprising after all and in line with previous research findings.

### Age influence on tyr - cognition relation

Next to sex/gender, also age seems to be relevant for the link between tyr in blood and executive attention. The positive association between serum tyr/LNAAs ratios and executive attention performance in the male sex/gender group was only significant when age was included as a control variable to account for negative age effects on task performance. We also showed that higher age was associated with higher dietary tyr intake. However, we were not able to confirm the interaction effects of sex/gender and age already described in previous research. Inconclusively, one study reported sex/gender effects (i.e. significant higher protein intake in male subjects) and sex/gender-age interactions with respect to dietary protein intake (Hone et al., 2020), whereas two other studies did not report any effect of age or sex/gender on dietary try (Hensel et al., 2019; Kühn et al., 2019). A longitudinal study showed increasing plasma tyr levels with increasing age (Darst et al., 2019) and another could demonstrate age differences in baseline serum tyr levels and in dose-dependent responses to oral tyr administration (van de Rest et al., 2017), yet both studies were conducted in aging populations. Contrary to previous findings, we could not identify any age effect or age-sex/gender interaction with respect to serum tyr levels, possibly due to the young to middle-aged sample.

### ANT performance

In contrast to previous studies (Gamboz et al., 2010; Jennings et al., 2007), we could not establish an effect of age on executive attention performance. Nevertheless, we found and interaction effect of age and sex/gender, revealing that the effect of age is only present in the male sex/gender group. Although previous studies found slower reaction times for executive function in older participants (mean age: 69.14/67.9), conflict effects did not differ significantly when controlling for general slowing in processing speed (Gamboz et al., 2010; Jennings et al., 2007). Interestingly, differences in executive attention performance (i.e. conflict effects RT) between younger and older adults were only found in the second experimental block of the ANT, suggesting a stronger influence of cognitive fatigue on older adults’ performance (Fu et al., 2021). However, this study did not control for general slowing at all.

In contrast, our analysis revealed no significant relationship between age and reaction times (although we did not even correct for general slowing with age), which is not surprising given the small age range and relatively young age of our sample. Importantly, our results on age effects on executive attention performance (in the male sex/gender group) should be interpreted with caution because we did not control for general slowing with age. This might be the main reason for reduced performance with higher age (Rey-Mermet & Gade, 2018) and should be included in future analyses.

Furthermore, the results of our whole-brain correlation analyses indicate a negative, locally non-specific association between age and white matter microstructure showing FA decreases with aging. This finding replicates the well-established age-related decline in white matter integrity (Cox et al., 2016; Vinke et al., 2018), starting already in young adulthood (Giorgio et al., 2010; Kodiweera et al., 2016).

### Limitations

Besides considerations for interpreting results mentioned above, there are limitations in the study design. First, meals consumed just before ANT may have influenced task performance by either stimulating effects (e.g., coffee) or digestion-related sedative effects (e.g., burgers). Second, indirect effects of AA content and macronutrient composition relating to lower serum tyr may have not been traceable due to the too short time interval between food intake and ANT. Indeed, previously it has been shown that although the plasma tyr/LNAAs ratios increase linearly after a meal, it takes up to two hours to reach highest levels (Strang et al., 2017), whereas in our design only 30 minutes had passed. In addition, additional time passes until tyr crosses the blood-brain barrier and is converted to DA; however, evidence is missing on how long transport and DA synthesis take in humans.

Next to confounding factors, differences in study procedure should be considered. For instance, testing sessions were scheduled at constant morning slots per participant, neglecting that serum tyr and tyr/LNAAs ratios vary throughout the day dependent on the amount of protein/tyr consumed (J D Fernstrom et al., 1979). Interestingly, serum tyr levels at 7 am were almost the same for all different diets and the tyr/LNAAs ratios of participants consuming 150 g and 75 g protein converged in the afternoon. The main limitation of the theoretical foundation concerns the reduction of relevant influencing factors and their interrelationships, which led to a simplistic and mechanistic model. For instance, with respect to dopaminergic neuromodulation, Cools points to three key principles (i.e., regional specialization, self-regulation, and baseline dependence) that indicate that the link between DA availability in the brain and cognitive function is not a one-way, monocausal, linear relationship (Cools, 2019). Moreover, it has been shown that chronically higher DA levels may lead to long-term adaptive changes in the dopaminergic system in terms of receptor or transporter expression and receptor sensitivity, which in turn may attenuate acute DA depletion effects (Hartmann et al., 2020). Also, brain DA in humans is mostly measured using proxy markers, such as precursor availability, and not directly using PET imaging, tracing techniques or manipulated by using dopaminergic drugs.

### Conclusions and outlook

In summary, non-pre-registered exploratory analyses revealed multiple sex/gender and age effects which are largely consistent with previous research, yet which might have possibly confounded the outcome of our analyses (Figure 14). Further exploration of sex/gender differences as imperfect proxies for yet unknown underlying biological and non-biological mechanisms is important, at least as a transitional solution to ensure equality in medical treatment and basic research for different sex/gender groups. Future neuroscientific research should consider differences in biological or non-biological mechanisms related to sex/gender when designing nutrition-related studies.

**Figure 13:**
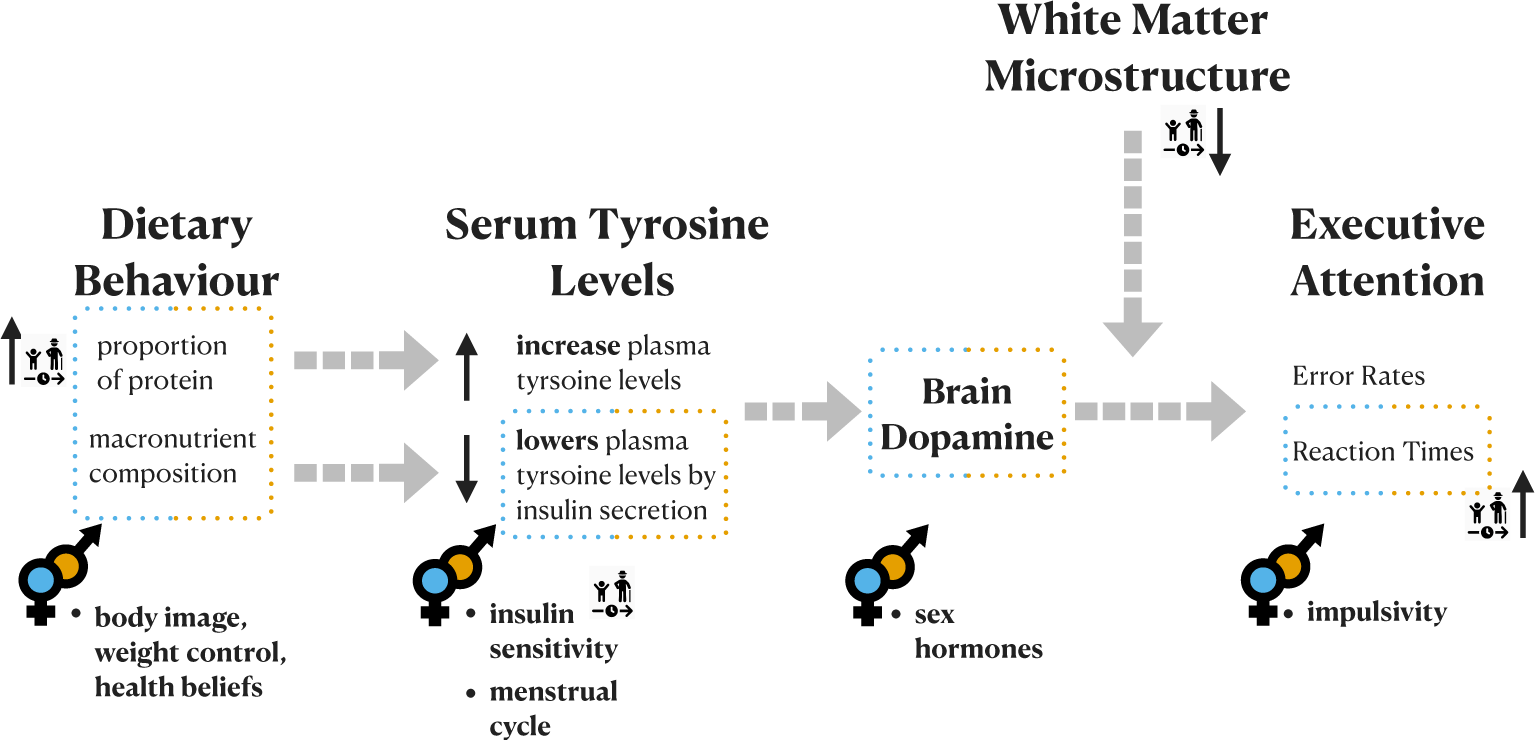
Aspects of sex/gender differences and age effects regarding the link between nutrition and cognition.

**Figure 14:**
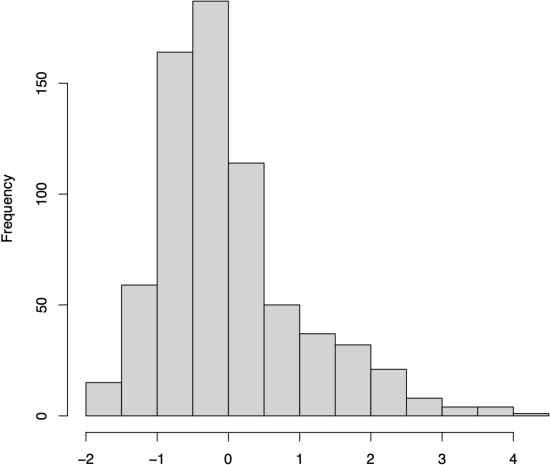
Histogram of standardized residuals (reaction times)

### Ethical statement

This study was performed in compliance with the Helsinki Declaration guidelines. The institutional ethics board of the Medical Faculty of the University of Leipzig, Germany, raised no concerns regarding the study protocol (228/18-ek) and all participants provided written informed consent. They received reimbursement for participation.

## Author contributions

Conceptualization, AKB, AVW, EM; methodology, AKB, AVW, EM, FB, RT; formal analysis, AKB; statistical advice: FB; data curation, AKB; data visualization, AKB; writing—original draft preparation, AKB; writing—review and editing, AVW, EM, RT; project administration, AVW, AV, JS. All authors have read and agreed to the published version of the manuscript.

## Acknowledgements

We thank all individuals for participating in the study and the medical staff for collecting blood samples. Also we thank Hendrik Hartmann for providing insights on considerations in dopaminergic markers and H. Lina Schaare for supporting data analysis regarding the Attention Network Test.

## List of abbreviations

executive attention: Executive Attention
FA: Fractional Anisotropy
DTI: Diffusion Tensor Imaging
DA: Dopamine
ANT: Attention Network Theory
RT: Reaction Time
ER: Error Rate
ACC: Anterior Cingulate Cortex
VTA: Ventral Tegmental Area
LNAA: large neutral amino acids
BOLD: blood oxygen level dependent
OSF: Open Science Framework
COS: Center for Open Science
BMI: Body-Mass-Index
SES: Socio-Economic Status
MRI: Magnetic Resonance Imaging
FFQ: German Food Frequency Questionnaire
TR: Repetition time
TE: Echo Time
FOV: Field-of-view
DW-MRI: Diffusion-weighted magnetic resonance imaging
GRAPPA: GeneRalized Autocalibrating Partial Parallel Acquisition
FSL: FMRIB Software Library
TBSS: Tract-based spatial statistic
ANOVA: analysis of variance
TFCE: Threshold-Free Cluster Enhancement
FWE: Family wise error
SD: standard deviation
Tyr: tyr
AA: amino acids

## Appendix

### 1.1. Pre-registered Analyses

#### ANT: Analysis of Variance (ANOVA)

Performing and correctly interpreting the results of an ANOVA requires that the assumption of normality and homoscedasticity is met.

First, the normal distribution of the standardized residuals was visually examined by plotting a histogram of standardized residuals and the Quantile-Quantile Plot based on both RT and ER data. Although the histogram of the RT data exhibits a slight skewness (Figure 14) and the data deviate from the reference line at the right edge of the Q-Q plot (Figure 15), we conclude that the visualization of the residuals sufficiently supports the normality assumption.

**Figure 15:**
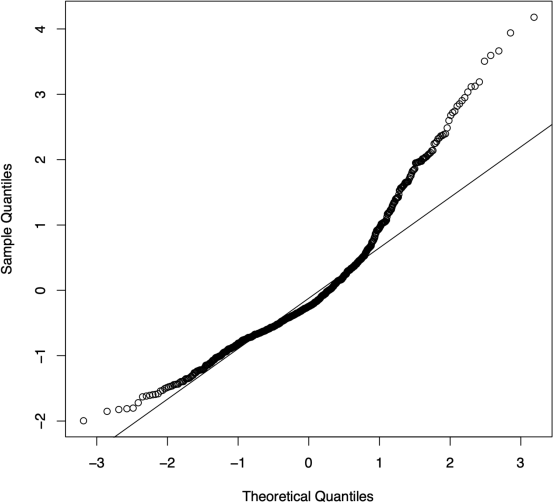
Quantile-Quantile Plot (reaction times)

However, the normality assumption is unambiguously violated with respect to the ER data (Figure 16 & Figure 17).

**Figure 16:**
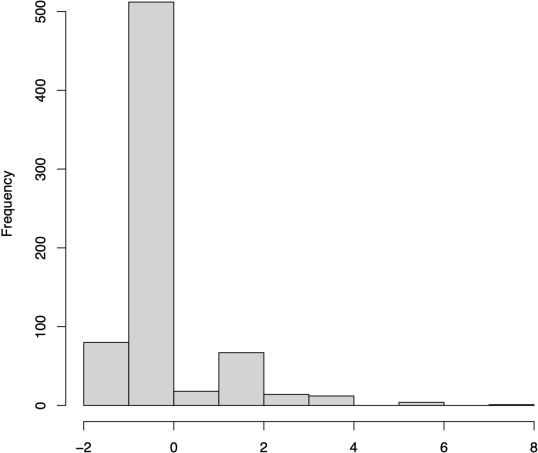
Histogram of standardized residuals (error rates)

**Figure 17:**
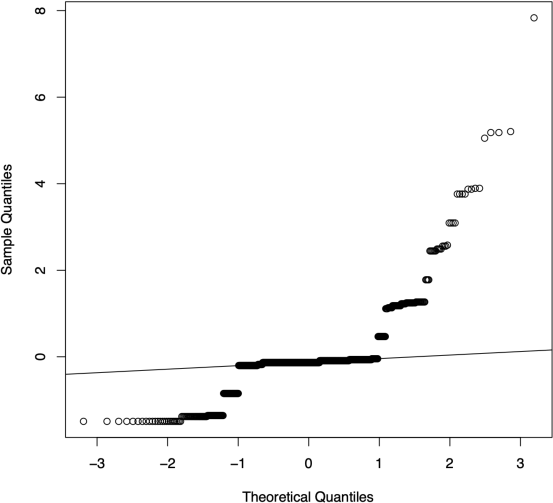
Quantile-Quantile Plot (error rates)

Second, the normality assumption was also tested analytically using the Shapiro-Wilk normality test (Shapiro & Wilk, 1965). The results for both RT (W = 0.92063, p-value < 2.2e-16) and ER (W = 0.70485, p-value < 2.2e-16) demonstrate that the null hypothesis (normality assumption) should be rejected. However, the results of analytical tests such as the Shapiro-Wilk normality test should be evaluated with caution, as even small deviations can lead to a rejection of the null hypothesis.

Third, homogeneity of variance was checked as well. Homoscedasticity concerns the assumption of equal or similar variances with respect to the different factors being compared. This assumption can be tested either visually or by Levene’s test (Levene, 1960). Regarding the RT data, both visual inspection (Figure 18) and Levene’s test (F(11,684) = 1.0253, p = 0.4217) confirms that the variance between factors is not significantly different i.e., the variances are homogeneous. However, regarding the ER data, both visual inspection (Figure 19) and Levene’s test (F(11, 696) = 27.698, p < 2.2e-16) indicate a lack of homogeneity; the p-value is smaller than the significance level of 0.05. Thus, the second assumption for a two-way ANOVA of the ER data could not be met either.

**Figure 18:**
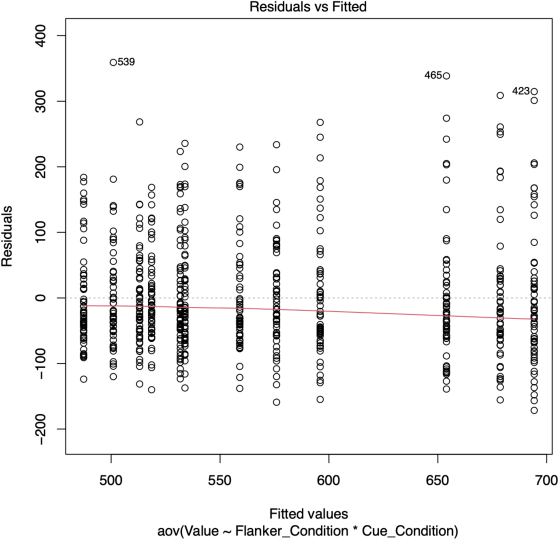
Residuals vs. Fitted plot (reaction times)

**Figure 19:**
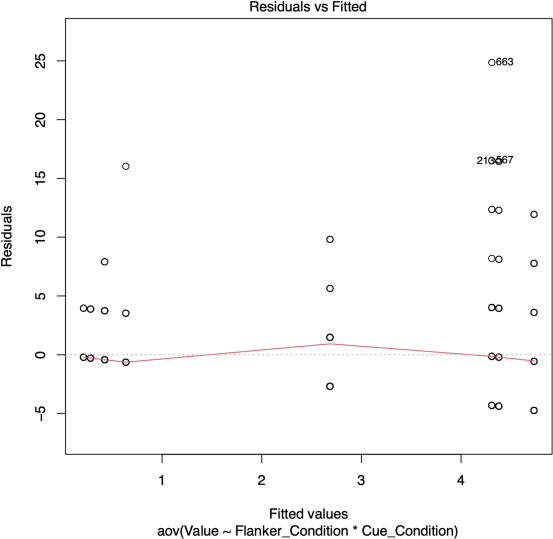
Residuals vs. Fitted plot (error rates)

#### Dietary - Serum Tyrosine Relationship

**Figure 20:**
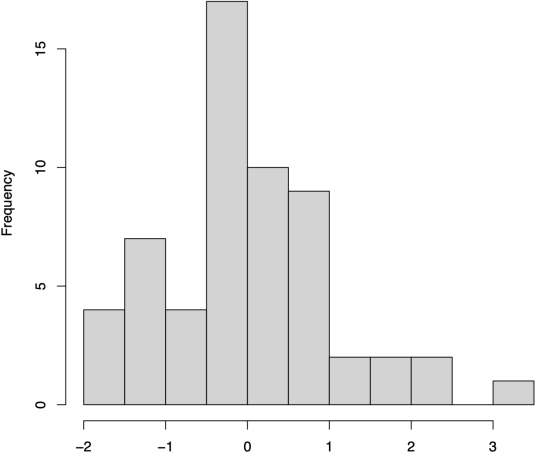
Histogram of standardized residuals: Dietary Tyrosine - Serum Tyrosine

**Figure 21:**
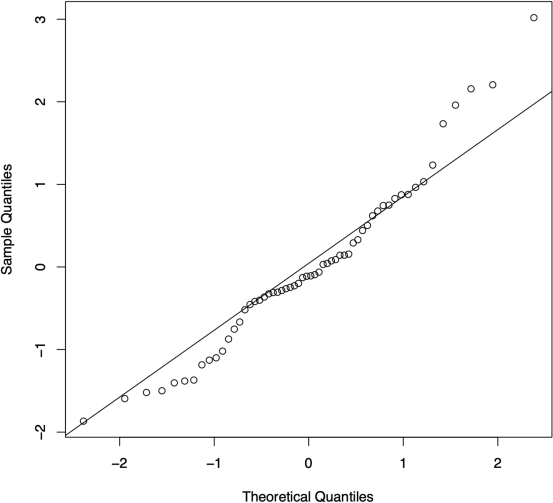
Normal Quantile-Quantile Plot: Dietary Tyrosine - Serum Tyrosine

Serum Tyrosine Ratio and Executive Attention Performance

**Figure 22:**
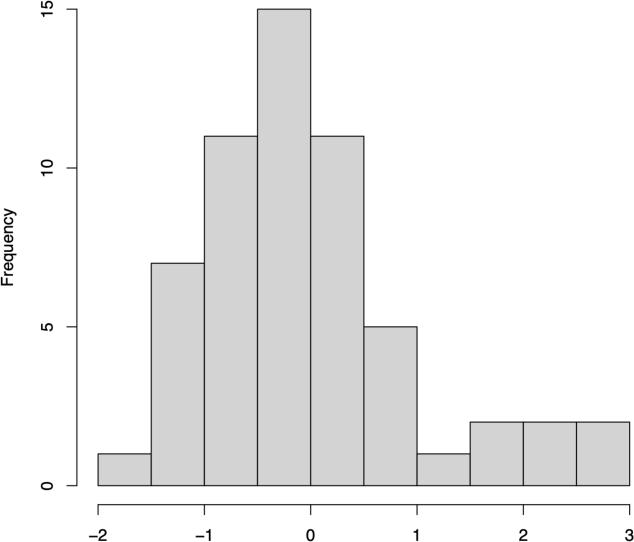
Histogram of standardized residuals: Serum Tyrosine/LNAA - EA Performance. Abbreviations: LNAA, large neutral amino acids; EA, executive attention.

**Figure 23:**
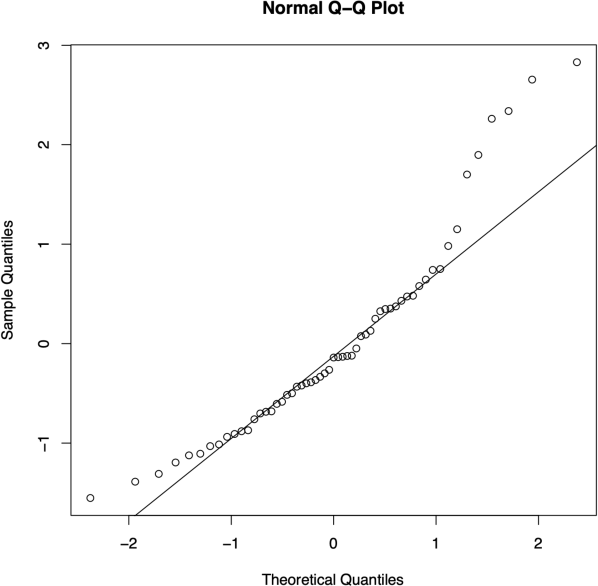
Normal Quantile-Quantile Plot: Serum Tyrosine/LNAA - EA Performance. Abbreviations: LNAA, large neutral amino acids; EA, executive attention.

#### Dietary Tyrosine Intake and Executive Attention

**Figure 24:**
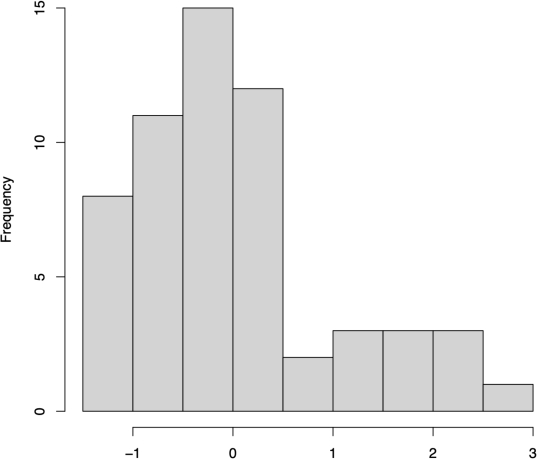
Histogram of standardized residuals: Dietary TYR/body weight - EA performance. Abbreviations: TYR, tyrosine; EA, executive attention.

**Figure 25:**
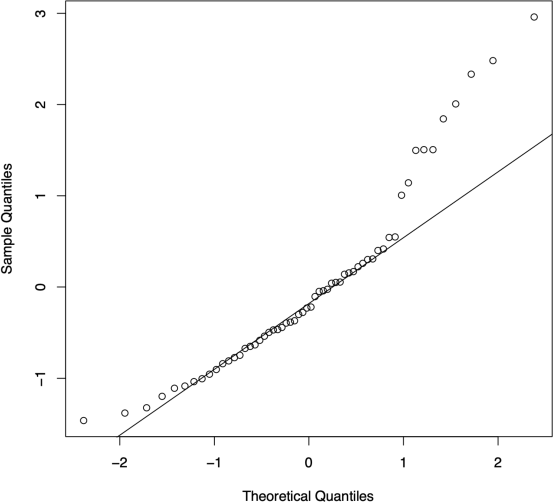
Normal Quantile-Quantile Plot: Dietary TYR/body weigh - EA performance. Abbreviations: TYR, tyrosine; EA, executive attention.

**Figure 26:**
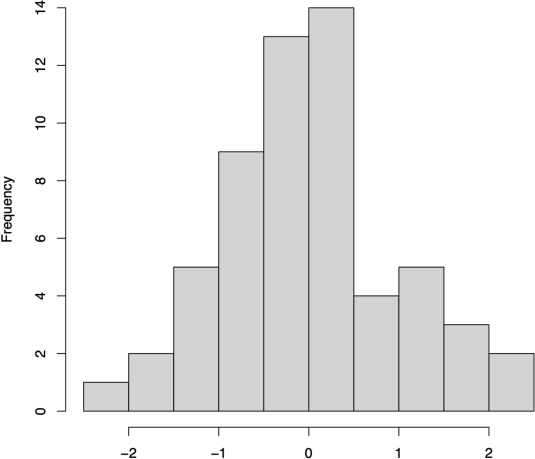
Histogram of standardized residuals: Dietary TYR/body weigh - EA performance (log-transformed data). Abbreviations: TYR, tyrosine; EA, executive attention.

**Figure 27:**
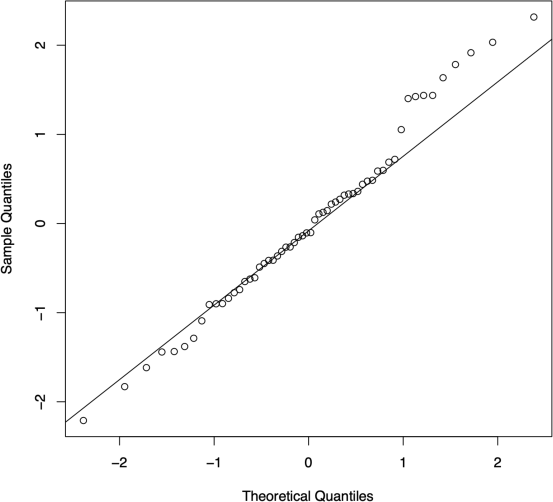
Normal Quantile-Quantile Plot: Dietary TYR/body weigh - EA performance (log-transformed data). Abbreviations: TYR, tyrosine; EA, executive attention.

### 1.2. Non-pre-registered Analyses

#### Sex/gender Differences in Variables of Interest

We plotted the histograms of the variable of interest. The visualization shows that the normality assumption is not met for almost all variables.

**Figure 28:**
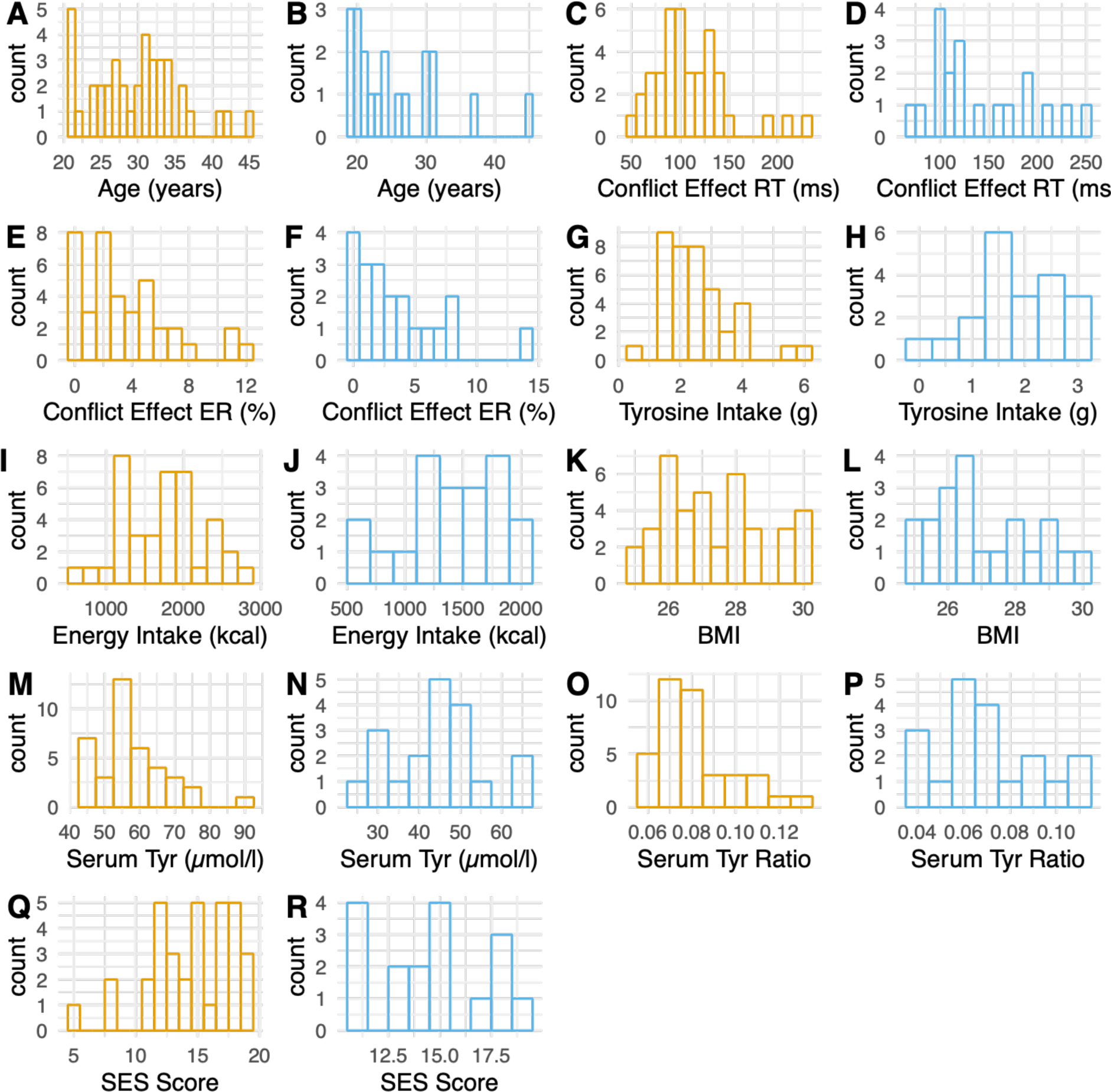
Histograms of variables of interest (blue: female sex/gender group, orange: male sex/gender group). Abbreviations: BMI, body mass index; TYR, tyrosine; SES, socio-economic status.

#### Regression Analysis per sex/gender

##### Female sex/gender group

**Figure 29:**
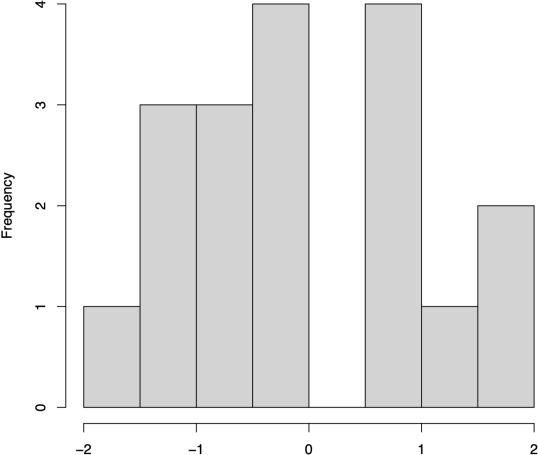
Histogram of standardized residuals: Serum tyrosine/LNAA ratio– EA (female*). Abbreviations: LNAA, large neutral amino acids; EA, executive attention.

**Figure 30:**
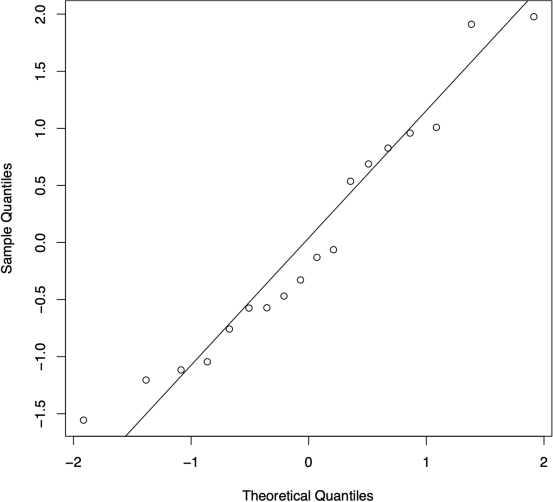
Normal Quantile-Quantile Plot: Serum tyrosine/LNAA ratio - EA (female*). Abbreviations: LNAA, large neutral amino acids; EA, executive attention.

##### Male sex/gender group

With regard to the significant multiple regression model examining whether serum tyr ratios significantly predicted executive attention performance in the male sex/gender group (when controlling for age), the normal distribution assumption was checked. First, the normal distribution of the standardized residuals was examined graphically. Both, the histogram and the Quantile-Quantile-Plot sufficiently support the normality assumption of the residuals; besides deviations on the outer edges. Secondly, the normality assumption was tested analytically using the Shapiro-Wilk normality test. The results also support that the null hypothesis of normal distribution cannot be rejected (W = 0.97719, p-value = 0.6021).

**Figure 31:**
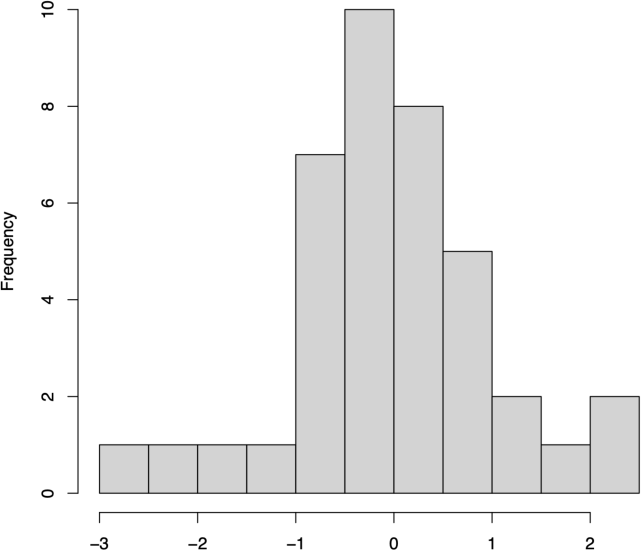
Histogram of standardized residuals: Serum tyrosine/LNAA ratio - EA (male*). Abbreviations: LNAA, large neutral amino acids; EA, executive attention.

**Figure 32:**
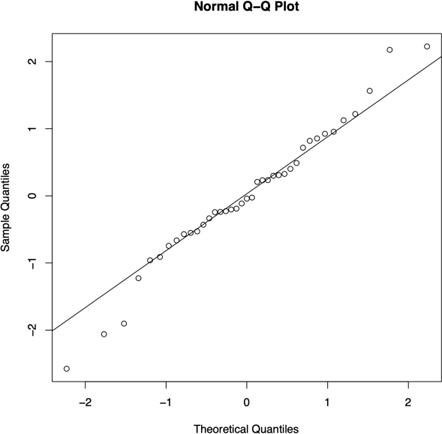
Normal Quantile-Quantile Plot: Serum tyrosine/LNAA ratio - EA (male*). Abbreviations: LNAA, large neutral amino acids; EA, executive attention.

##### Age-Relationships

Checking Normality assumption visually and by analytical test:

Regression model: Age - executive attention (W = 0.90476, p-value = 0.0002519) – no normal distribution

Regression Model: Age - Diet (W = 0.94139, p-value = 0.00683) – no normal distribution

Regression Model: Age - Serum tyr (W = 0.98677, p-value = 0.7789) - normal distribution

**Figure 33:**
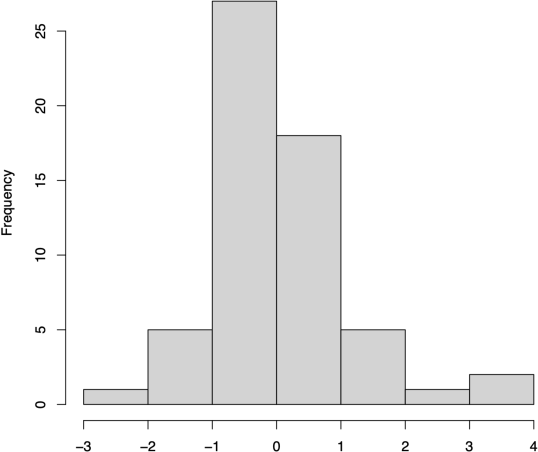
Histogram of standardized residuals: Age - Dietary TYR. Abbreviations: TYR, tyrosine.

**Figure 34:**
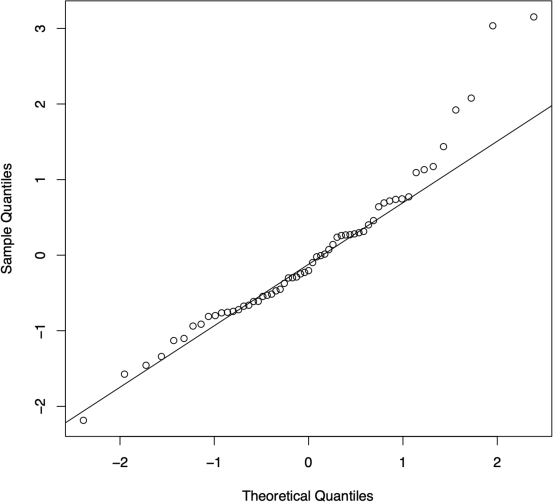
Normal Quantile-Quantile Plot: Age -Dietary TYR intake. Abbreviations: TYR, tyrosine.

**Figure 35:**
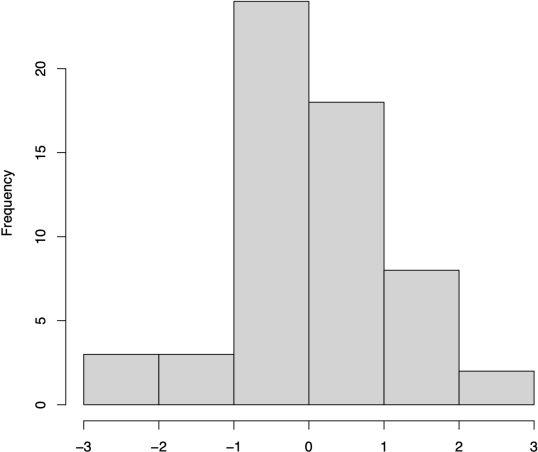
Histogram of standardized residuals: Age - Serum TYR levels. Abbreviations: TYR, tyrosine.

**Figure 36:**
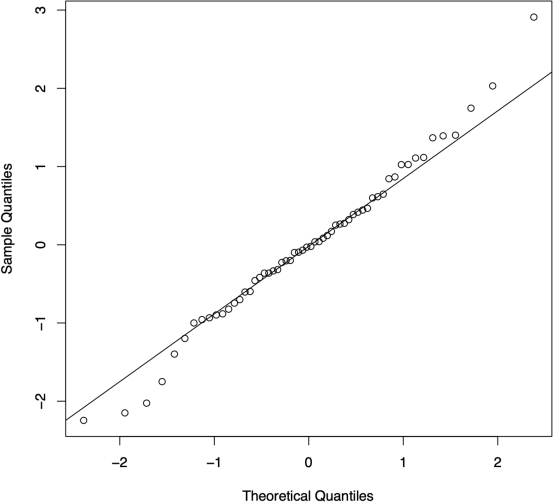
Normal Quantile-Quantile Plot: Age - Serum Tyrosine levels.

**Figure 37:**
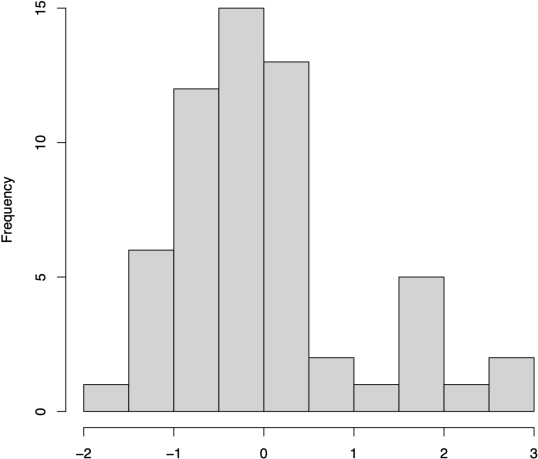
Histogram of standardized residuals: Age - EA performance. Abbreviations: EA, executive attention.

**Figure 38:**
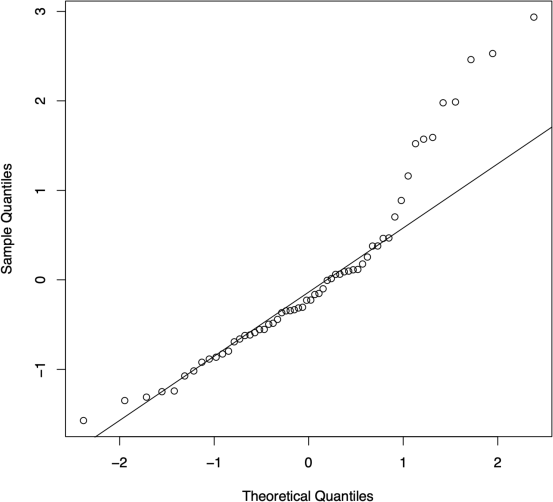
Normal Quantile-Quantile Plot: Age - Executive Attention Performance

##### Age: sex/gender interaction - executive attention performance

**Figure 39:**
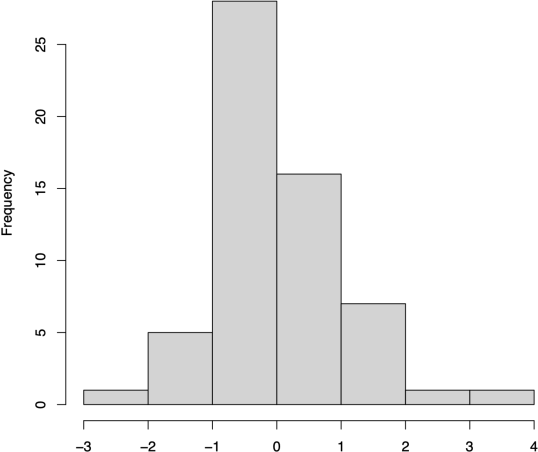
Histogram of standardized residuals: Diet TYR - Age:Sex/gender. Abbreviations: TYR, tyrosine.

**Figure 40:**
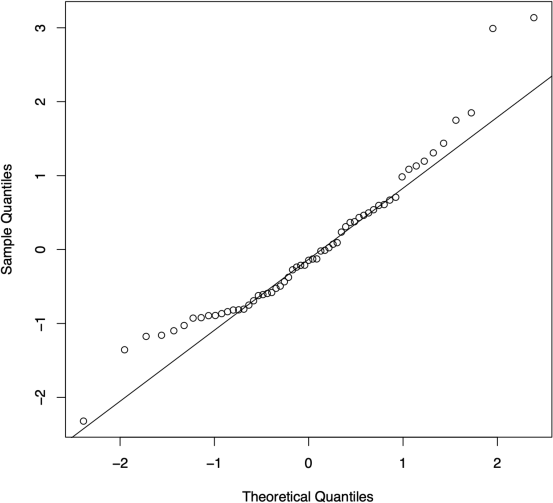
Normal Quantile-Quantile Plot: Diet TYR - Age:Sex/gender. Abbreviations: TYR, tyrosine.

**Figure 41:**
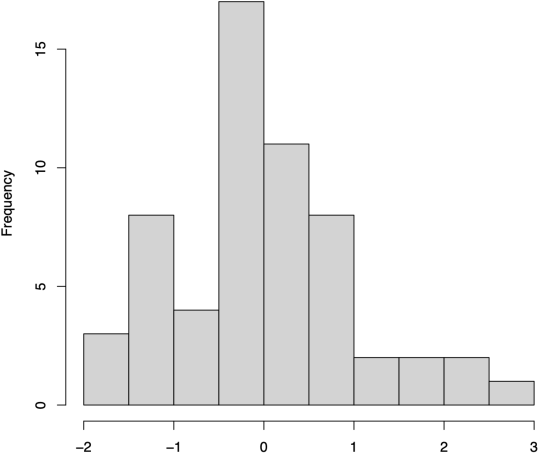
Histogram of standardized residuals: Serum TYR - Age:Sex/gender. Abbreviations: TYR, tyrosine.

**Figure 42:**
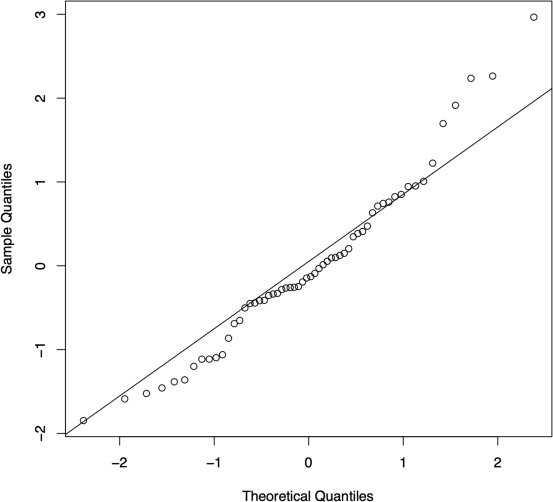
Normal Quantile-Quantile Plot: Serum TYR - Age:Sex/gender. Abbreviations: TYR, tyrosine.

**Figure 43:**
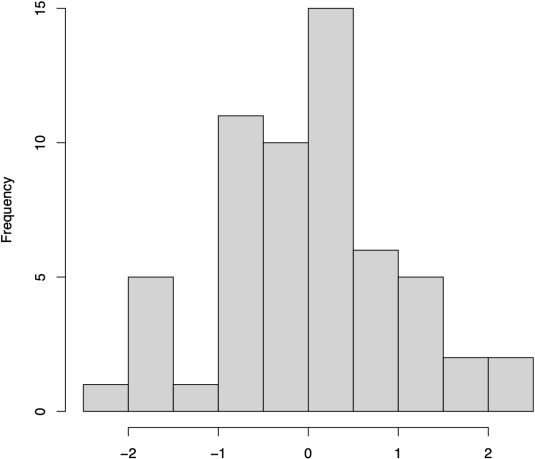
Histogram of standardized residuals: EA - Age:Sex/gender (log-transformed data). Abbreviations: EA, executive attention.

**Figure 44:**
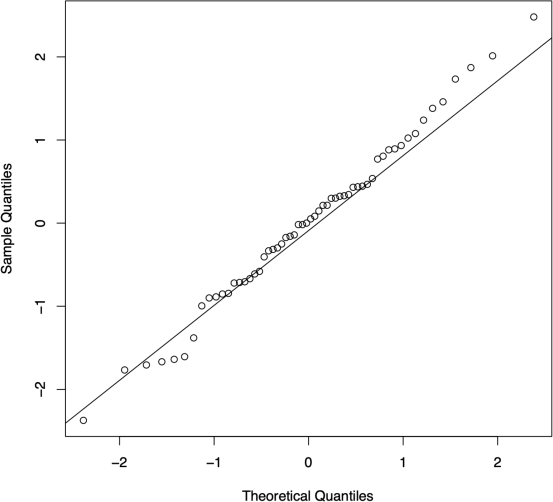
Normal Quantile-Quantile Plot: EA - Age:Sex/gender (log-transformed data). Abbreviations: EA, executive attention.

##### Conflict Effect ER

Conflict Effect computed based on Error Rate data. We performed non-parametric Mann-Whitney U/Wilcoxon rank sum tests. The results show no significant sex/gender difference in executive attention performance based on error rate data (W = 391.5, p-value = 0.9807).

**Figure 45:**
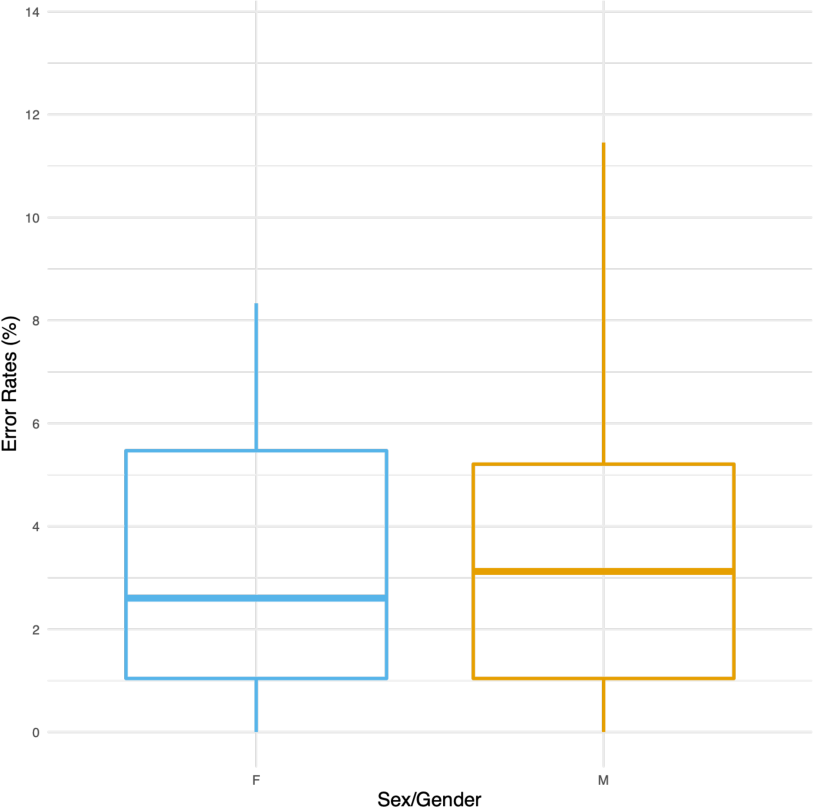
Sex/Gender group difference in conflict effects ER (%). Abbreviations: ER, error rates.

**Table 9:**
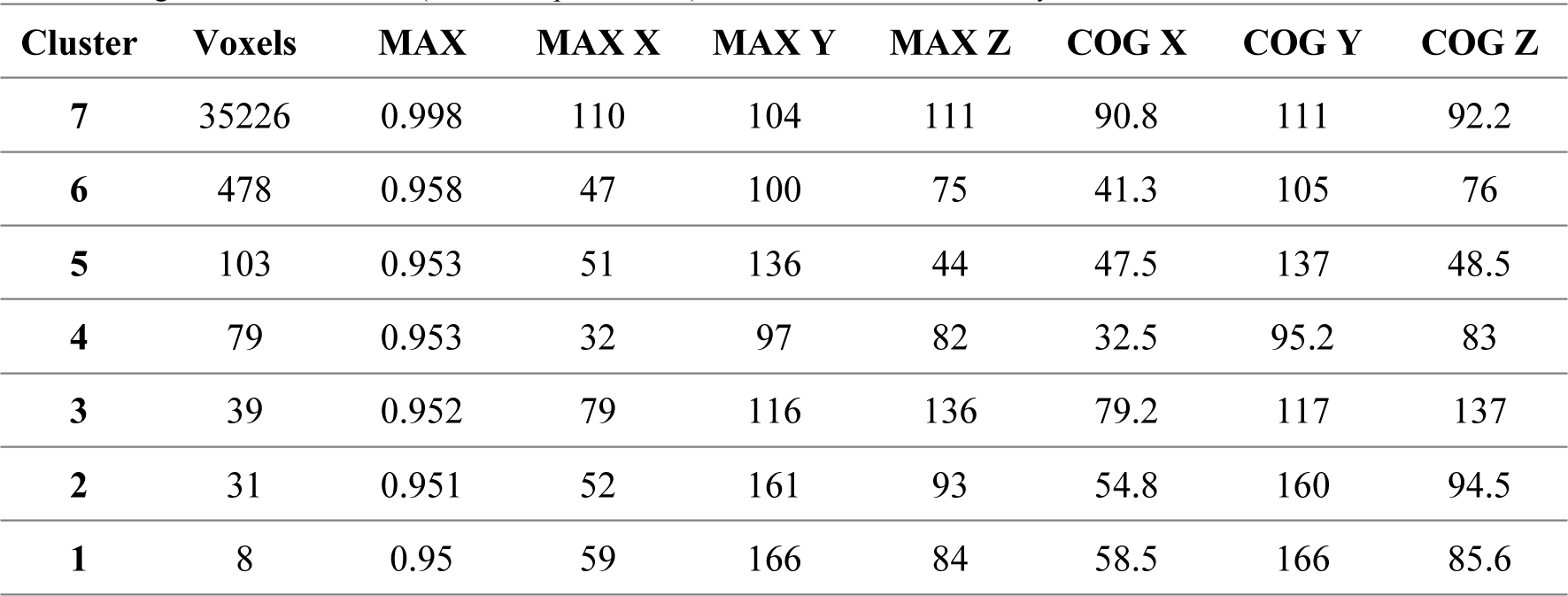
7 Age-associated clusters (threshold: p_FWE_ < 0.05). Abbreviations: FWE, family-wise error corrected.

**Table 10:**
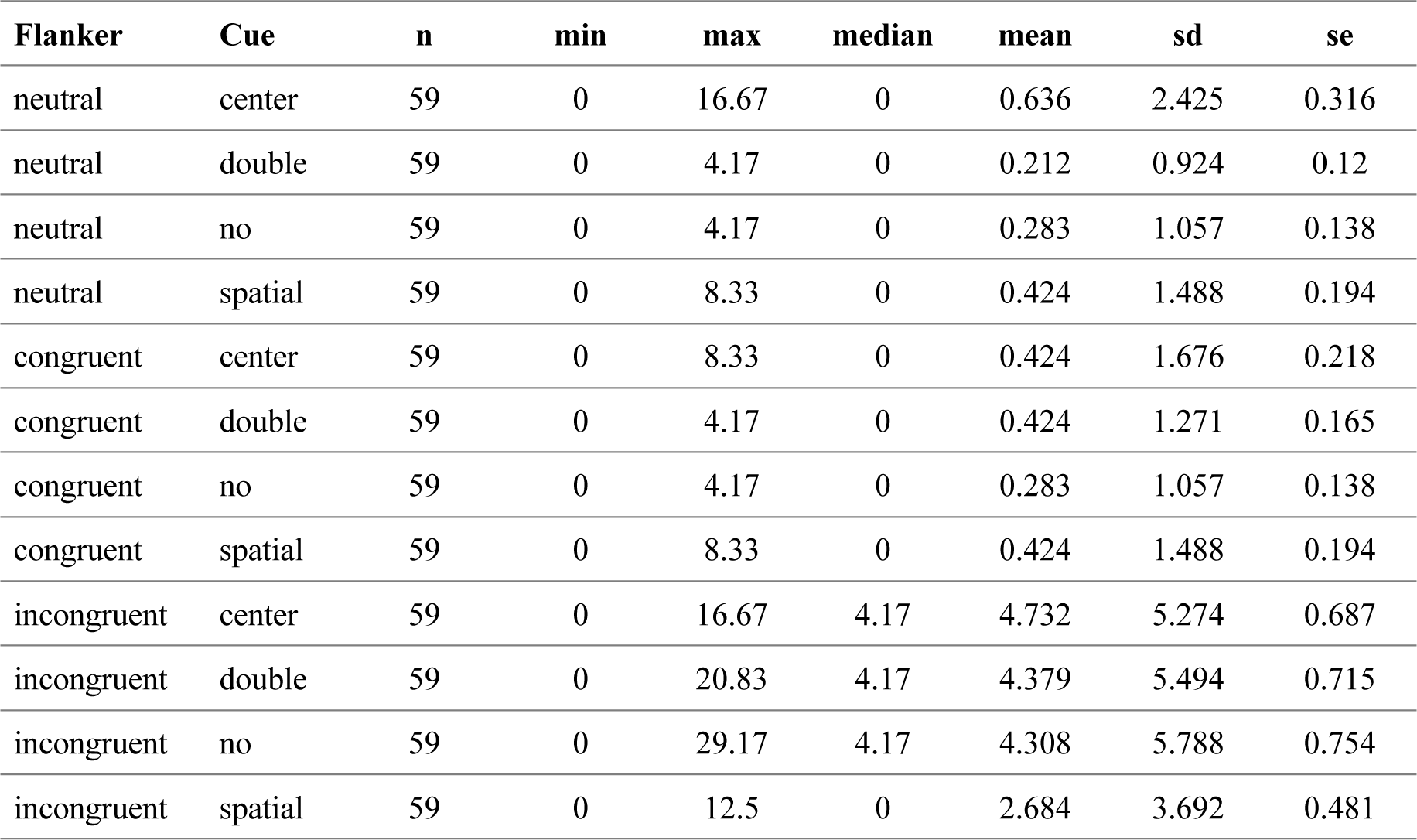
Summary statistics of error rates as a function of cue condition and flanker type (in %). Abbreviations: n, number of subjects; sd, standard deviation; se, standard error.

**Table 11:**
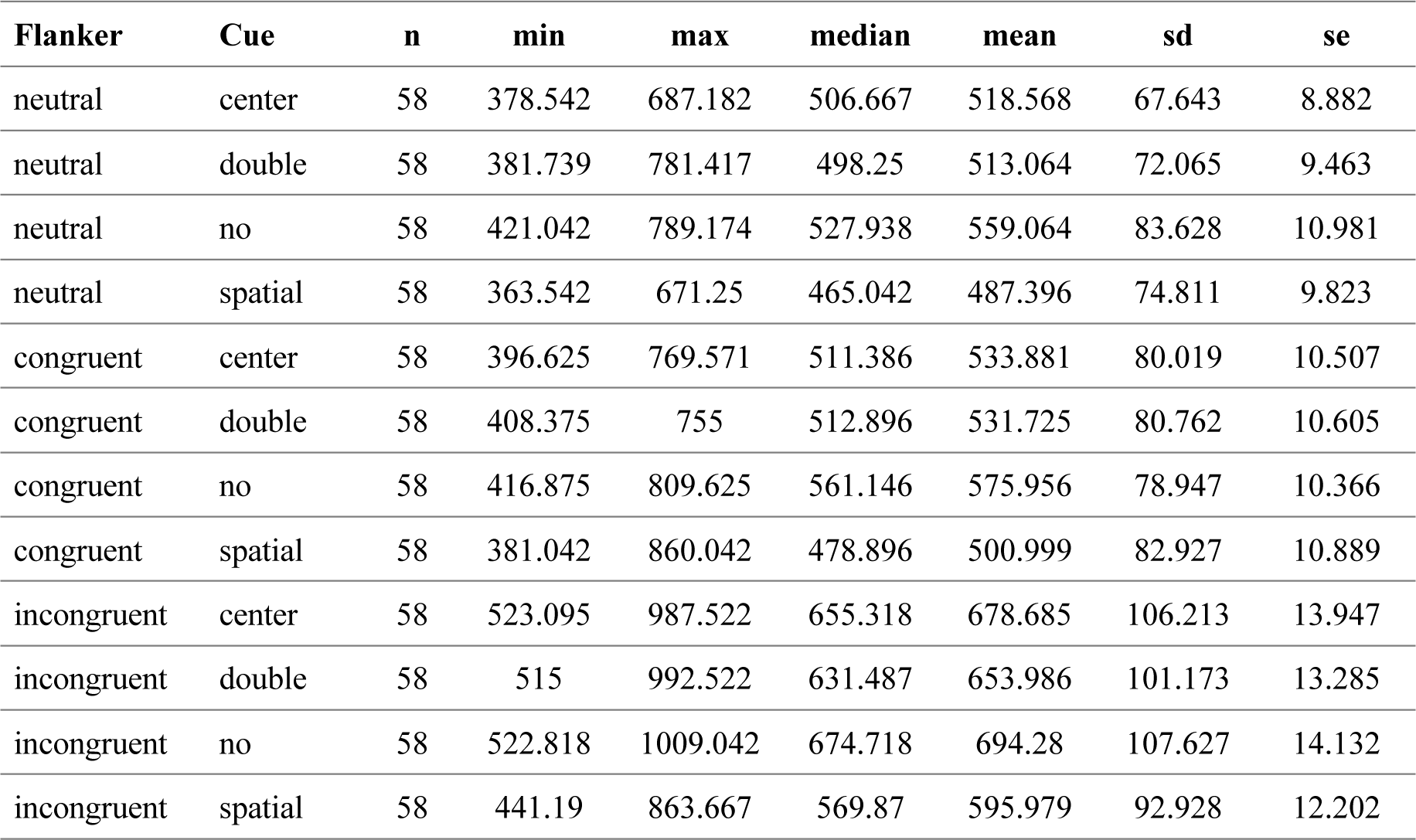
Summary statistics of reaction times as a function of cue condition and flanker type (in ms). Abbreviations: n, number of subjects; sd, standard deviation; se, standard error.

